# Ultrabright NIR-II Nanoprobes for *Ex Vivo* Bioimaging: Protein Nanoengineering Meets Molecular Engineering

**DOI:** 10.1101/2025.07.27.665782

**Authors:** Isabella Vasquez, Asma Harun, Robert Posey, Ruhan Reddy, Nikita Gill, Ulrich Bickel, Joshua Tropp, Indrajit Srivastava

## Abstract

Near-infrared (NIR) fluorescence imaging is a powerful, non-invasive tool for cancer diagnosis, enabling real-time, high-resolution visualization of biological systems. While most probes target the first NIR window (NIR-I, 750-950 nm), recent advances focus on the second window (NIR-II, 1000-1700 nm), which offers deeper tissue penetration and reduced interferences from scattering and autofluorescence. However, many current NIR-II nanoprobes show suboptimal brightness and limited validations in more human-centric models. Here, we present an orthogonal strategy combining molecular engineering, by modulating the amount and position of thiophene moieties in semiconducting polymers (SPs), with protein nanoengineering to develop ultrabright NIR-II imaging probes optimized for *ex vivo* bioimaging in large animal models. The molecular tuning amplifies the NIR-II fluorescence brightness while screening endogenous proteins as encapsulating matrices to improve colloidal stability and enable active targeting. Molecular docking identified bovine serum albumin as the effective candidate, and the resulting protein-complexed nanoprobes were characterized for size, colloidal stability under physiological conditions, and optical performances. Imaging performances were evaluated using tumor-mimicking phantoms in porcine lungs, simulating cancer surgery, and injected at clinically relevant concentrations into ovine brains and porcine ovaries for microvascular visualization and tissue discrimination, respectively. In all scenarios, our protein-complexed nanoprobes outperformed the FDA-approved clinical dye indocyanine green in signal-to-background ratios. Initial *in vitro* assays confirmed their hemocompatibility, biocompatibility, and cellular uptake in ovarian adenocarcinoma cells. This integrated approach offers a promising platform for developing next-generation ultrabright NIR-II nanoprobes with improved brightness and stability, advancing the potential for image-guided surgery and future clinical translation.

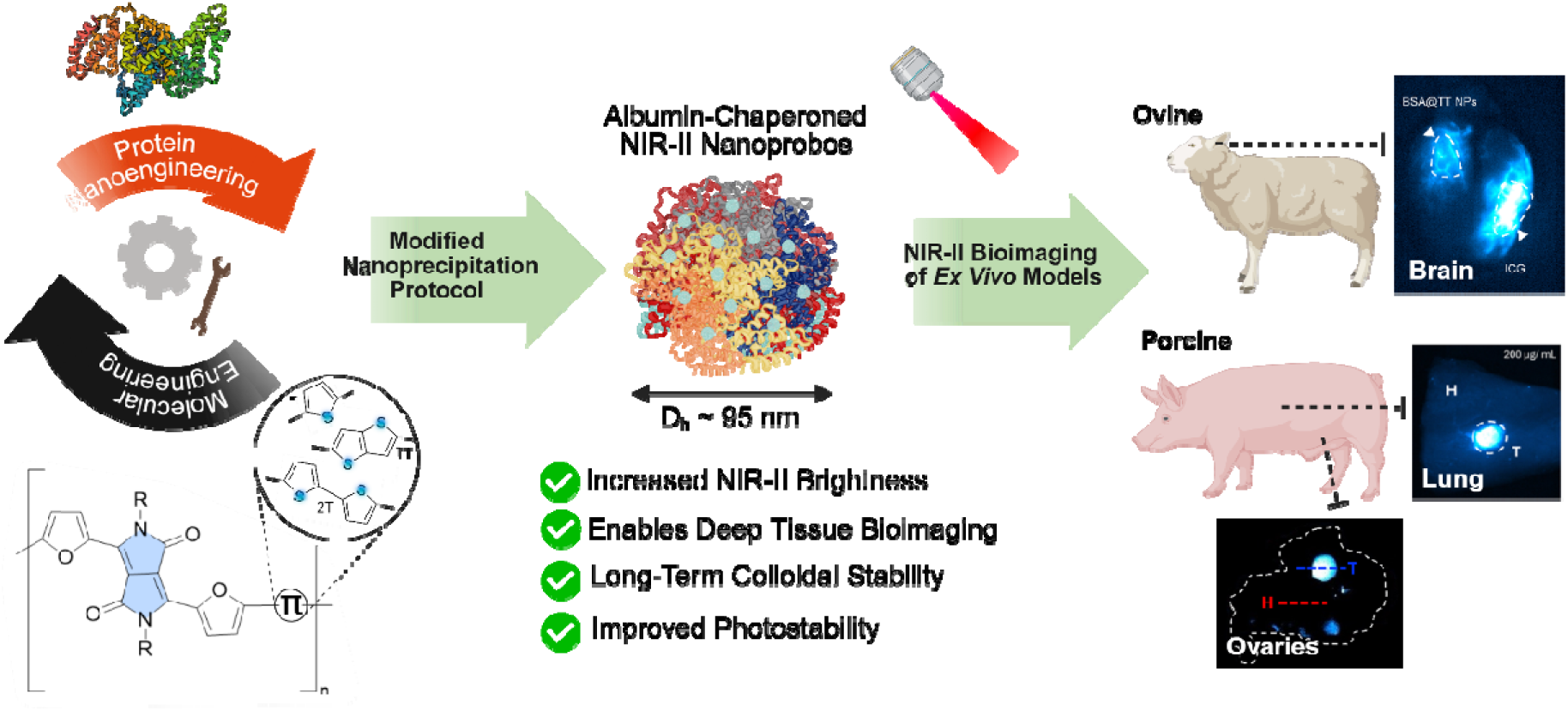

## INTRODUCTION

Light-based techniques have revolutionized biomedical imaging and diagnostics in recent years. ^1–3^ Near-infrared (NIR) fluorescence imaging has had a substantial impact on clinical diagnosis and navigation surgery owing to its operational simplicity and high spatial resolution.^4–6^ However, despite its exuberance, high sensitivity, real-time monitoring, and non-invasive features, its reliance on the visible or near-infrared (NIR-I, 700-900 nm) region suffers from poor tissue penetration and limited spatial resolution. Ushering in a new generation of NIR imaging in the second near-infrared (NIR-II, 1000-1700 nm) window has presented a promising solution preceding the traditional visible light or NIR-I window, whereby it significantly reduces tissue scattering and autofluorescence. These ideal optical traits improve penetration depth and impart high signal-to-background ratios (SBR) with excellent resolution of the anatomic structures due to the intrinsic advantages of negligible light scattering from tissues.^7–15^

The pursuit of NIR-II fluorescence imaging and the clinical need for improved biological imaging has led to a significant interest in the development of NIR-II probes capable of emitting in the NIR-II window. To date, varieties of NIR-II probes, including single-walled carbon nanotubes, quantum dots, metal nanoclusters, and semiconducting polymers (SPs), have been developed as promising candidates for constructing probes for pre-clinical bioimaging applications in the NIR-II spectral window.^16–24^ Among them, nanoprobes developed from semiconducting polymers have shown the most significant potential in terms of clinical applications due to their minimal toxicity concerns and impressive optical properties, including absorption cross-section, photostability, and fluorescence quantum yield; however, they do require further improvements.^25^ To this end, molecular engineering strategies have been actively pursued to enhance the performance of SPs. These strategies include tuning donor–acceptor– donor (D–A–D) molecular frameworks, modifying polymer backbones, or engineering side chains to systematically investigate and improve their optical performance, including NIR-II brightness.^24, 26–30^

Building on recent advancements, semiconducting polymers (SPs), which are typically hydrophobic and water-insoluble, generally require processing with amphiphilic polymers to form hydrophilic nanoparticles known as semiconducting polymer nanoparticles (SPNs), enabling their use in *in vitro*, *in vivo*, and *ex vivo* bioimaging applications.^36–39^ (Table 1) A commonly used amphiphilic matrix is DSPE-PEG2000, which enhances the colloidal stability and aqueous dispersibility of SPNs.^40, 41^ However, while DSPE-PEG2000 improves nanoparticle dispersion, it does not significantly enhance key optical properties such as fluorescence brightness, particularly in the NIR-II window.^37, 42^ As a result, the brightness of SPNs remain limited, restricting their effectiveness in larger animal models despite promising performance in small-animal studies.^43–45^

**Table 1.** A comparative table with important optical properties of recent NIR-II probes for varying medical imaging applications, including IRT^31^, Flav7^32, 33^, IR-E1^34^, H1^35^, and BSA@TT.

| NIR-II Probe | Solubility | Absorption/<br>Emission<br>(808 nm Exc.) | QY (%) | Targeting Moiety/<br>Application |
| --- | --- | --- | --- | --- |
| IRT | Water- soluble | 750/1050 nm | ~1.5% (Ref. dye: IR26)<sup>31</sup> | Targets tumors via the CD133 biomarker |
| Flav7 | Water- insoluble/ water-<br>soluble with lipid coating | 1026/1045 nm | 0.61% (Ref. dye: IR26)<br><sup>32,33</sup> | Blood and lymphatic<br>vessel imaging |
| IR-E1 | Water-soluble with lipid<br>coating | 830/1071 nm | ~0.7% (Ref. dye: IR26)<br><sup>34</sup> | Traumatic brain injury<br>imaging |
| H1 | Water- insoluble/ water-<br>soluble with lipid coating | 840/1100 nm | ~2.0% (Ref. dye: IR26)<br><sup>35</sup> | Tumor vascularization and<br>whole-body imaging |
| BSA@TT | Water- soluble | 672/960 nm | 3.82% (Ref. dye: ICG) | Tumor and<br>microvasculature imaging |

This limitation underscores the need for alternative amphiphilic materials that offer functional benefits beyond stabilization. One such candidate is albumin, a naturally abundant, biocompatible protein with intrinsic binding capabilities.^46–48^ Recent studies have shown that albumin can enhance both the fluorescence brightness and photostability of cyanine dyes, largely due to its ability to encapsulate these dyes within its hydrophobic pockets through covalent interactions.^49^ This encapsulation reduces nonradiative decay pathways caused by intramolecular rotations or vibrations, thereby significantly increasing the fluorescence quantum yield. Additionally, the albumin shell protects against collisional quenching and contributes to improved photostability and biocompatibility.^50–52^ Albumin-based carriers may also enable active targeting via gp60-mediated endocytosis.^53, 54^ While albumin encapsulation has shown promise for small-molecule dyes, systematic comparisons between SPNs stabilized with DSPE-PEG2000 *via* conventional nanoengineering and those coated with albumin through protein-based nanoengineering remain limited. Thus, the direct construction of albumin-coated SPNs represents a practical and promising strategy that is currently underexplored and lacks a head-to-head comparison with conventional amphiphilic formulations. Moreover, to our knowledge, no study has directly investigated the combined use of molecular engineering and protein nanoengineering strategies in SPNs to assess their respective and synergistic impacts on enhancing NIR-II optical performance.

Here, in this work, we adopt a dual approach to develop high-performance NIR-II SPN-based imaging nanoprobes by combining molecular engineering with protein nanoengineering strategies. Specifically, we tuned the SP backbone by varying the number and placement of thiophene-based donors within the D-A backbone to improve fluorescence brightness in the NIR-II window. The three backbone variations used in this study are a single thiophene unit (T), thienothiophene (TT), and 2-2’-bithiophene (2T) (Figure 1). In parallel, we explored protein nanoengineering to replace conventional amphiphilic coatings. Through molecular docking, bovine serum albumin (BSA) was identified as a highly compatible encapsulation matrix, offering both stability and biological functionality (Figure 1). The resulting BSA-coated SPN nanoprobes were systematically characterized for size, charge, colloidal behavior, and optical performance under physiological conditions. Their imaging capabilities were tested in realistic biological models, including tumor phantoms in porcine lungs and clinically relevant animal systems such as ovine brains and porcine ovaries, demonstrating superior visualization of microvasculature and tissue boundaries, respectively. Across all experiments, our new nanoprobes outperformed indocyanine green (ICG), a Food and Drug Administration (FDA)-approved dye, in signal-to-background ratios. Biocompatibility was validated through preliminary *in vitro* toxicity assays, and cellular uptake studies demonstrated enhanced internalization in tumor cells. Overall, this combined engineering strategy provides a promising route for advancing the design of ultrabright, stable NIR-II nanoprobes suitable for image-guided interventions and potential clinical applications.

**Figure 1.**
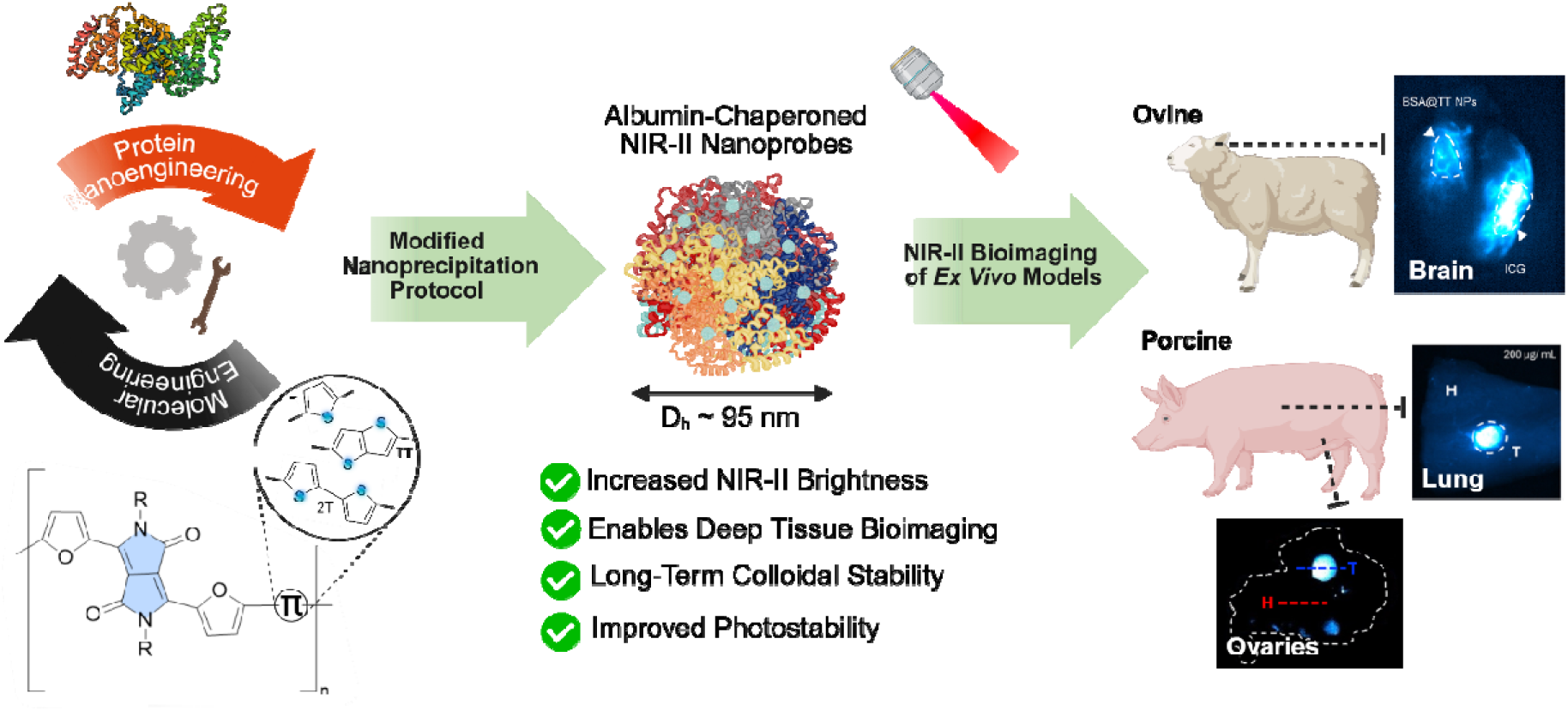
Schematic illustration of bovine serum albumin (BSA)-coated semiconducting polymers with varying thiophene donors within the D-A backbones (T, TT, and 2T). After dye encapsulation, the resulting nanoprobes (BSA@T, TT, 2T) exhibit ultrabright NIR-II fluorescence and have been applied towards bioimaging applications in multiple *ex vivo* tissue models.

## MATERIALS AND METHODS

### Materials

All materials were purchased from Sigma-Aldrich and used without modification, unless otherwise stated. Porcine skin tissue was obtained from local grocery stores. Preserved ovine brains with dura mater, porcine ovaries, and dry-preserved porcine lung tissue were purchased from Nasco Education.

### Molecular Docking

The molecular docking between Bovine Serum Albumin (BSA) and the thiophene-derivative polymers was performed using the UCSF Chimera software. The 3D crystal structure of BSA (PDB ID: 4f5s) was chosen as the receptor for molecular docking. The ligand structure of the three SPs (PDFT-T, PDFT-2T, PDFT-TT) was generated using the Avogadro software. Docking simulations were carried out using a single repeat unit of the PDFT polymer as the ligand. This representative monomer was chosen because simulating full-length polymer chains is computationally prohibitive and beyond the scope of conventional docking, while a single repeat unit provides a practical approximation of local binding interactions with BSA.^55, 56^ Following Dock Prep preparation of the protein, specific parameters for the active pocket of the protein were defined for the grid box. PDFT-T fits within the dimensions x=68.49, y=25.40, z=92.10 and docked with dimensions x=87.70, y=57.28, z=87.83 for chain-B of BSA. PDFT-2T fits within the dimensions x=7.41, y=18.35, z=101.93, with docking dimensions of x=99.54, y=73.70, z=66.33 for chain-A of BSA. PDFT-TT fits within the dimensions x=65.71, y=28.11, z=91.30 and docks with the dimensions x=75.72, y=59.45, z=75.12 for chain-B of BSA. Using the docked protein complexes, the Pymol software was used to determine binding energies and clarify the interacting amino acids. In this study, the integrated AutoDock Vina engine was used to execute docking in UCSF Chimera. To estimate binding affinities, AutoDock Vina uses a stochastic global optimization technique that combines an empirical scoring system with a gradient-based conformational search. Chimera was utilized for docking posture analysis, binding area definition, and receptor and ligand production.

### Synthesis and Preparation of PDFT SPs

Polymers were synthesized using a previously reported procedure, including our recent report.^38, 57–59^ The reactive monomers were subjected to Stille polycondensation (Scheme S1) with commercial 2,5-bis(trimethylstannyl)thiophene, 2,5-bis(trimethylstannyl)thieno[3,2-b]thiophene, and 5,5’-bis(trimethylstannyl)-2,2’-bithiophene donors to afford alternating copolymers PDFT-T and PDFT-TT, and PDFT-2T, respectively. NMR data (Figures S1–S4) match prior reports. Preparation of monomer (**2**) in brief: 1.2 g (1.45 mmol) of 3,6-di(furan-2-yl)-2,5-bis(2-octyldodecyl)-2,5-dihydropyrrolo[3,4-c]pyrrole-1,4-dione (**1**) was treated with N-bromosuccinimide (2.0 equiv) in dry CH_2_Cl_2_ at rt for 24 hours. The crude product was dissolved in methanol, recrystallized, and washed with cold methanol to give 3,6-bis(5-bromofuran-2-yl)-2,5-bis(2-octyldodecyl)-2,5-dihydropyrrolo[3,4-c]pyrrole-1,4-dione (**2**). ^1^H NMR (400 MHz, CDCl_3_) δ 8.29 (d, J = 3.7 Hz, 2H), 6.61 (d, J = 3.7 Hz, 2H), 3.97 (d, J = 7.6 Hz, 4H), 1.52 (s, 2H), 1.32 – 1.19 (m, 64H), 0.85 (dd, J = 6.6 Hz, 12H).

### Preparation of DSPE-PEG@T

For preparing DSPE-PEG@T, 0.5 mg of T polymer was co-dissolved in 2 mL of THF with 5 mg of surfactant, DSPE-PEG2K. This was followed by a dropwise addition to 8 mL PBS1X solution in a 20 mL vial under probe sonication for 4 min. THF was subsequently evaporated from the solution by stirring at 450 rpm at room temperature overnight. The mixture solution was ultrafiltered at 4500 rpm for 10 min and washed 3 times with PBS1X to remove any excess precursors. The nanoparticle solution was filtered using a 0.22-micron PTFE (Polytetrafluoroethylene) filter, and the final product (DSPE-PEG@T) was collected and stored at 4 °C.

### Preparation of DSPE-PEG@2T

For preparing DSPE-PEG@2T, 0.5 mg of 2T polymer was co-dissolved in 2 mL of THF with 5 mg of surfactant, DSPE-PEG2K. This was followed by a dropwise addition to 8 mL PBS1X solution in a 20 mL vial under probe sonication for 4 min. THF was subsequently evaporated from the solution by stirring at 450 rpm at room temperature overnight. The mixture solution was ultrafiltered at 4500 rpm for 10 min and washed 3 times with PBS1X to remove any excess precursors. The nanoparticle solution was filtered using a 0.22-micron PTFE (Polytetrafluoroethylene) filter, and the final product (DSPE-PEG@2T) was collected and stored at 4 °C.

### Preparation of DSPE-PEG@TT

For preparing DSPE-PEG@TT, 0.5 mg of TT polymer was co-dissolved in 2 mL of THF with 5 mg of surfactant, DSPE-PEG2K. This was followed by a dropwise addition to 8 mL PBS1X solution in a 20 mL vial under probe sonication for 4 min. THF was subsequently evaporated from the solution by stirring at 450 rpm at room temperature overnight. The mixture solution was ultrafiltered at 4500 rpm for 10 min and washed 3 times with PBS1X to remove any excess precursors. The nanoparticle solution was filtered using a 0.22-micron PTFE (Polytetrafluoroethylene) filter, and the final product (DSPE-PEG@TT) was collected and stored at 4 °C.

### Preparation of BSA@T

0.5 mg of T polymer was dissolved in 2 mL of THF. 52.8 mg of BSA was dissolved in 8 mL of PBS1X in a separate vial and heated to approximately 37 °C to ensure homogeneity. The THF solution was then added dropwise to 8 mL of BSA-PBS1X solution and sonicated for 4 min. THF was subsequently evaporated overnight by stirring at 450 rpm. The mixed solution was ultrafiltered at 4500 rpm for 10 min and washed 3 times with PBS1X to remove any excess precursors or unreacted BSA. The nanoparticle solution was filtered using a 0.22-micron PTFE (Polytetrafluoroethylene) filter, and the final product (BSA@T) was collected and stored at 4 °C.

### Preparation of BSA@2T

0.5 mg of 2T polymer was dissolved in 2 mL of THF. 52.8 mg of BSA was dissolved in 8 mL of PBS1X in a separate vial and heated to approximately 37 °C to ensure homogeneity. The THF solution was then added dropwise to 8 mL of BSA-PBS1X solution and sonicated for 4 min. THF was subsequently evaporated overnight by stirring at 450 rpm. The mixed solution was ultrafiltered at 4500 rpm for 10 min and washed 3 times with PBS1X to remove any excess precursors or unreacted BSA. The nanoparticle solution was filtered using a 0.22-micron PTFE (Polytetrafluoroethylene) filter, and the final product (BSA@2T) was collected and stored at 4 °C.

### Preparation of BSA@TT

0.5 mg of TT polymer was dissolved in 2 mL of THF. 52.8 mg of BSA was dissolved in 8 mL of PBS1X in a separate vial and heated to approximately 37 °C to ensure homogeneity. The THF solution was then added dropwise to 8 mL of BSA-PBS1X solution and sonicated for 4 min. THF was subsequently evaporated overnight by stirring at 450 rpm. The mixed solution was ultrafiltered at 4500 rpm for 10 min and washed 3 times with PBS1X to remove any excess precursors or unreacted BSA. The nanoparticle solution was filtered using a 0.22-micron PTFE (Polytetrafluoroethylene) filter, and the final product (BSA@TT) was collected and stored at 4 °C.

### Physicochemical and Optical Characterization

Hydrodynamic diameter and polydispersity index (PDI) were determined via dynamic light scattering (DLS), using the Litesizer DLS 700 Particle Analyzer instrument (Anton Paar). Measurements were recorded using solutions of 30 µL of nanoparticle solution combined with 970 µL of PBX1X. ζ-potential measurements also used the Litesizer DLS 700 Particle Analyzer. Ultraviolet-visible-near infrared (UV-VIS-NIR) absorption was recorded using the Genesys 30 Visible Spectrophotometer (ThermoFisher Scientific). All absorbance measurements were recorded with scanning intervals of 1 nm from 325 to 1100 nm. Fluorescence characterization was performed using a dual absorbance and fluorescence spectrophotometer (Olis DSM 142 UV/Vis/NIR) featuring extended InGaAs detectors to assess the NIR emission of each SP. Fluorescence spectra were obtained by exciting at the corresponding λ_max, abs_ of each SP.

### IR Vivo Imaging of NPs

Microcentrifuge tubes with stock solutions of NPs at concentrations of 200 µg/mL were imaged with the IR Vivo imager (Photon etc.). Fluorescence images were collected with the excitation wavelengths 760 and 808 nm and 3 emission filters: NIR-I (800 – 975 nm), NIR-II (1000 – 1600 nm), and LP1250 (1250-1600 nm).

### Evaluating Fluorescence Quenching in Blood

100 µL of BSA@TT (200 µg/mL) was mixed with 100 µL of 10% packed porcine red blood cells in a microcentrifuge tube at room temperature. Using the IR Vivo imager (Photon etc.), images were collected for excitation wavelength 808 nm and emission windows NIR-I, NIR-II, and LP1250. The packed porcine red blood cells used in this study were obtained from Innovative Research.

### Relative Quantum Yield Calculations

Relative quantum yield () was determined using ICG in PBS1X as the reference standard (taken as = 1.7%).^60^ The fluorescent intensity at 808 nm excitation was plotted against absorbance for four different concentrations of each nanoprobe, where absorbance was < 0.1, and the slope from the linear fit was used in the following equation:

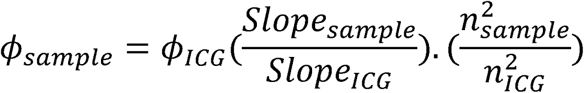

where *_sample_* and *_ICG_* are the quantum yields of the sample and reference, respectively. Slope_sample_ and Slope_ICG_ are the slopes of the NIR-II fluorescent intensity vs. absorbance plots obtained from the four concentrations, and *n* is the refractive index (equal to 1.334 for all samples in PBS1X).

### Extinction Coefficient Determination

For each nanoprobe/contrast agent: ICG, DSPE-PEG@TT, and BSA@TT, absorbance measurements were collected at four different concentrations where the absorbance was <0.1. A linear regression of absorbance at 808 nm versus concentration was used to determine ε_m_ in units of L ^-1-1^

### Fluorescence Brightness Calculation

Fluorescence brightness was calculated as the product of the mass extinction coefficient and quantum yield:

Brightness = ε_m_ ×

Brightness values were calculated for each nanoprobe using the ε_m_ and values derived from the datasets.

### Fluorescence titration assay for BSA-TT binding

Fluorescence titration experiments were performed to evaluate the binding affinity of TT with BSA. A fixed concentration of TT (1000 μM; prepared in PBS, pH 7.4) was incubated with increasing concentrations of BSA (0–100 μM) at room temperature for 30 min to allow equilibrium binding. After addition, each mixture was briefly vortexed (5–10 s) and gently shaken to ensure homogeneous complex formation. Fluorescence spectra were collected using a plate reader (λ_ex_. = 785 nm; λ_em_. = 900 nm), and the maximum emission intensity was recorded for each titration point. Background signals from buffer-only wells were subtracted. The resulting binding curve was plotted as fluorescence intensity (a.u.) versus BSA concentration (μM). Data were fitted to a one-site binding hyperbola model using GraphPad Prism 9.0 to obtain the equilibrium dissociation constant (K_d_) and maximum binding capacity (B_max_). All experiments were performed in triplicate, and data are presented as mean ± SD.

### Encapsulation efficiency of BSA@TT nanoprobes

Encapsulation efficiency was determined by UV–Vis spectroscopy. A calibration curve was constructed using TT solutions in THF (10– 200 µg/mL), with absorbance at the maximum wavelength plotted against concentration. The absorbance spectrum of the BSA@TT nanoprobes was then collected under the same conditions. Using the calibration curve, the concentration of TT within the BSA@TT formulation was calculated and compared to the initial TT amount used in the formulation. Encapsulation efficiency was expressed as the percentage of TT incorporated into the nanoprobes relative to the starting amount.

### *Ex Vivo* Porcine Tissue Penetration Studies

To evaluate the optical performance of BSA@TT through varying tissue depths, NMR tubes containing 1 mL of BSA@TT solution (200 µg/mL) were imaged through increasing porcine skin thicknesses (0, 2, and 4 mm) with the IR Vivo imager (Photon etc.). Images were collected for excitation wavelengths of 760 and 808 nm and emission windows of NIR-I, NIR-II, and LP1250. Imaging exposure time was altered according to laser wavelength, emission window, and penetration depth. The orientation of the tubes does not affect the fluorescence measurements, as all samples were imaged under identical excitation and collection conditions.

### *Ex Vivo* Ovine Brain and Porcine Ovary Experiment

Sheep brains with dura mater and pig ovaries were obtained from NASCOguard. 200 µL of ICG (10 µM) and 200 µL of BSA@TT (200 µg/mL) were injected approximately 2 cm below the surface of the superior frontal lobe and posterior parietal lobe, respectively. To avoid significant fluorescence interference, injections were done on laterally opposite regions of the brains (n=3). 100 µl of ICG (10 µM) and 100 µL of BSA@TT (200 µg/mL) were injected directly into the follicles of separate porcine ovaries (n=3). The IR Vivo imager (Photon etc.) was used to collect fluorescent images with an 808 nm laser and emission windows ranging from 850 to 1600 nm.

### *Ex Vivo* Porcine Lung Tumor Delineation Experiment

Dry-preserved swine lungs were obtained from SPECTRUM Nasco Educational Supplies. Tumor-mimicking phantoms with increasing concentration of BSA@TT (10, 50, 100, and 200 µg/mL) and ICG (10 µM) were created using a recipe previously reported by our group (n=2).^61, 62^ To simulate an exposed tumor lesion following surgical resection, each tumor phantom was imaged in a hollowed portion of the lung using the IR Vivo imager (Photon etc.). Images were collected for excitation wavelengths of 760 and 808 nm and emission windows of NIR-I, NIR-II, and LP1250.

### Cytotoxicity Evaluation in Ovarian Cancer Cells

To assess the cytotoxicity effects of DSPE-PEG@TT and BSA@TT NPs after 48 hours, a colorimetric MTT assay was used for both treatments at increasing concentrations (1, 5, 10, 25, 50, and 100 µg/mL). A control group was established by only treating cells with reconstituted Dulbecco’s Modified Eagle Medium (DMEM). OVCAR8 cells were plated overnight in a 96-well plate at 10^5^ cells per well, and each well was treated with 20 µL of a 5mg/ mL MTT solution.^63–66^ After 4 hours of incubation at 37°C and 5% CO_2_ in the Heracell Vios 160i (Thermo Scientific), the formazan crystals were dissolved with 200 µL of DMSO. The DMSO-filled wells were incubated for an additional 15 minutes at 37°C and shaken for 5 seconds. Immediately following this, the absorbance values were collected at 592 nm using the BioTek Epoch 2 Microplate Spectrophotometer (Agilent).

## RESULTS AND DISCUSSION

### Design and Synthesis of PDFT Copolymers

To achieve absorption or emission in the NIR within semiconducting conjugated polymers (SPs), donor-acceptor (D-A) backbones are typically required to narrow the bandgap through molecular orbital hybridization and perturbation theory. ^24, 67^ Due to the energy gap law, very few organic materials feature bright emission in the NIR-II – also referred to as the shortwave infrared (SWIR).^68^ As there are few examples of D-A copolymers that emit strongly beyond 1000 nm, our approach centers on modifying a previously reported, high-performing SP (PDFT) featuring the furan-flanked diketopyrrolopyrrole acceptor unit.^38^ We recently showed that adjusting the electron-donating thiophene unit within the PDFT backbone is a viable strategy for tuning the polymer’s fluorescence brightness.^58, 69, 70^ In particular, the copolymer featuring a thieno[3,4-b]thiophene (TT) donor exhibited superior performance, which is attributed to the increased planarity and rigidity of its polymer backbone.

Here, we selected three electron-rich donor motifs, thiophene (T), bithiophene (2T), and thieno[3,4-b]thiophene (TT), to introduce increasing degrees of π-conjugation and backbone rigidity (Figure 2a). These donors provide a rational design space for exploring how conformational flexibility affects both photophysical properties and protein binding. More flexible backbones (e.g., PDFT-2T) may adopt different orientations within albumin’s hydrophobic pockets compared to more rigid analogs like PDFT-TT, which we previously showed enhances NIR-II brightness due to increased planarity and reduced torsional disorder.^71, 72^ These structural differences could directly influence encapsulation behavior, nanoprobe stability, and bioimaging performance. Each of the PDFT materials was synthesized according to previous reports using microwave-assisted Stille copolymerization procedures.^57, 58^ The weight-average molecular weight (*M*_w_) and dispersity (Đ) were determined by high-temperature gel permeation chromatography (HT-GPC) at 120 °C in 1,2,4-trichlorobenzene (1 mg mL ¹, stabilized with 125 ppm of BHT) using a Tosoh EcoSEC High Temperature GPC system with two TSK gel columns in series (G3000Hhr by G2000Hhr) at a flow rate of 1 mL min ¹ (Figure 2b, 2c). The optical properties of the polymers were then measured, confirming that their absorption and emission spectra were consistent with prior reports (Figure 2d, 2e).

**Figure 2.**
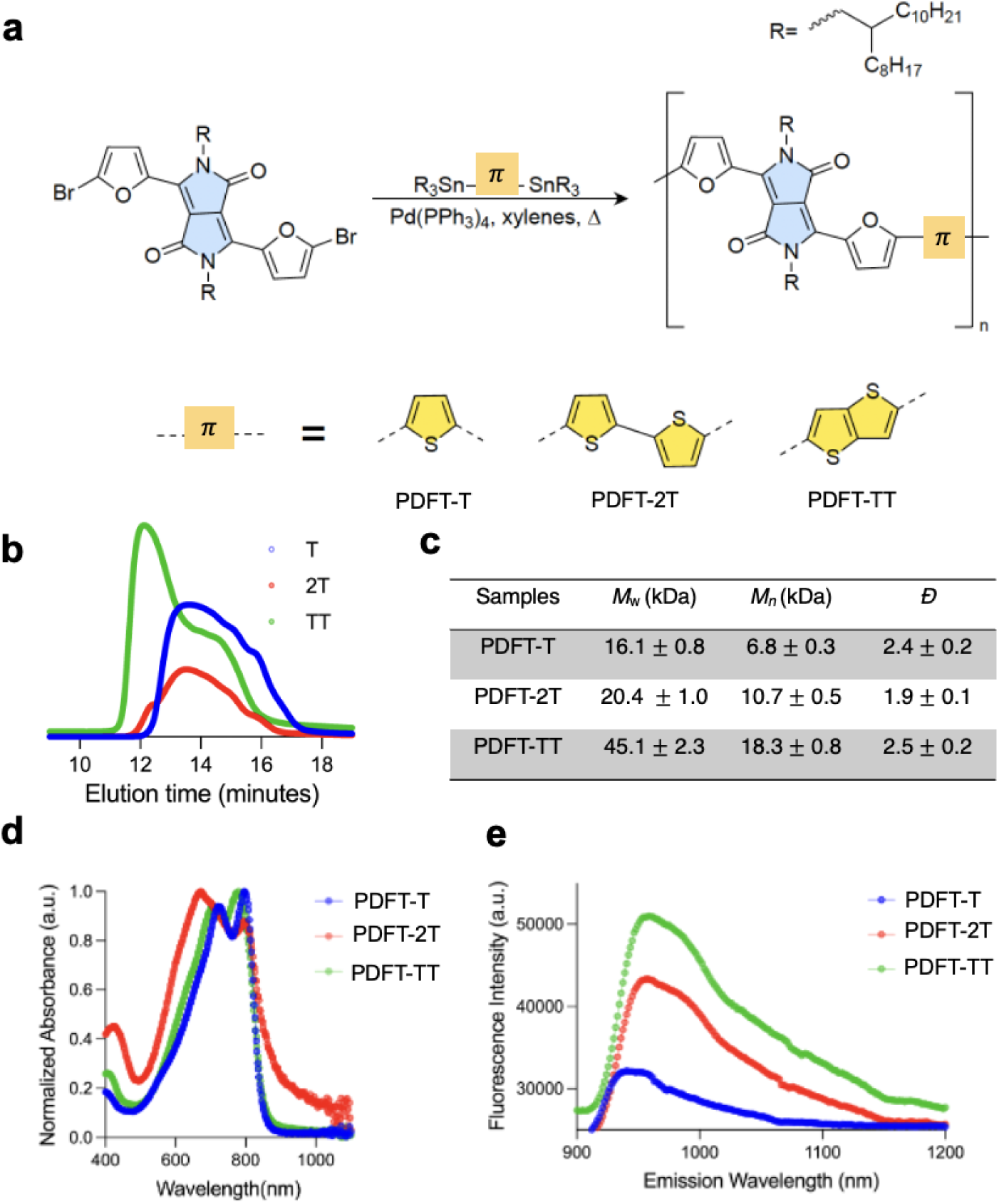
Design and Synthesis of PDFT Copolymers. (a) Polymerization of PDFT-T, PDFT-2T, and PDFT-TT copolymers with branched aliphatic side chains via Stille polycondensation. (b) High-temperature gel-permeation chromatography (GPC) of each polymer batch with (c) corresponding structural analysis. (d) Normalized absorption and (e) representative near infrared emission optical profiles of each D-A SP in THF.

### *In Silico* Analysis of Protein Interactions with T-Derivative Polymers

To evaluate the theoretical binding sites of the thiophene-derived polymers (T, TT, and 2T) investigated in this study, we conducted a series of computational modeling experiments. These analyses aimed to (i) identify the specific protein chains associated with each polymer, and (ii) visualize the amino acid residues involved in each binding pocket, along with their interactions with various segments of the polymer structures (Figure 3a-g). The binding energies varied according to the polymers’ attachment sites: -6.1 kcal/mol for PDFT-T (or T); -6.4 kcal/mol for PDFT-2T (or 2T); and -7.3 kcal/mol for PDFT-TT (TT) (Figure 3h). The reported docking binding energy (– 7.3 kcal/mol) represents the overall Gibbs free energy change of the protein–polymer complex and is consistent with non-covalent interactions such as hydrogen bonds, hydrophobic contacts, van der Waals forces, and electrostatics.^73^ These favorable interactions are partly counterbalanced by entropic penalties, desolvation effects, and conformational adjustments required for binding.^74^ The resulting moderate binding energy is therefore fully consistent with reversible noncovalent complexation, as further supported by our experimental fluorescence titration studies (Kd ≈ 76 µM). PDFT-T and PDFT-TT docked within the B chain of BSA’s tertiary structure and interacted with the well-characterized Sudlow binding site I.^75^ Despite targeting the same site, their binding scores differed due to the variations in polar bonding, ligand orientation, and the surrounding hydrophobic amino acids.^75, 76^ Notably, PDFT-TT (TT)’s structure enables the formation of two covalent bonds with oxygen atoms in the polymer’s backbone, facilitating interactions with both negatively charged glutamate and positively charged arginine residues (Figure 3i). The difference in binding affinities among the polymers can also be attributed to the number of aromatic rings present in each backbone. Since Sudlow site I is known to preferentially bind aromatic compounds, such as ibuprofen and diflunisal, the additional thiophene ring in PDFT-TT’s structure may account for its stronger association with BSA at this particular location.^77, 78^ Interestingly, PDFT-2T preferentially binds to a hydrophobic pocket found in Domain III of BSA. Although PDFT-2T and PDFT-TT share similar overall aromaticity, the sigma bond separating the thiophene units in 2T resulted in markedly different binding orientation and affinity towards BSA. Overall, this preliminary docking study demonstrated that the optimal candidate for albumin chaperoning within these three thiophene variations is PDFT-TT (or TT).

**Figure 3.**
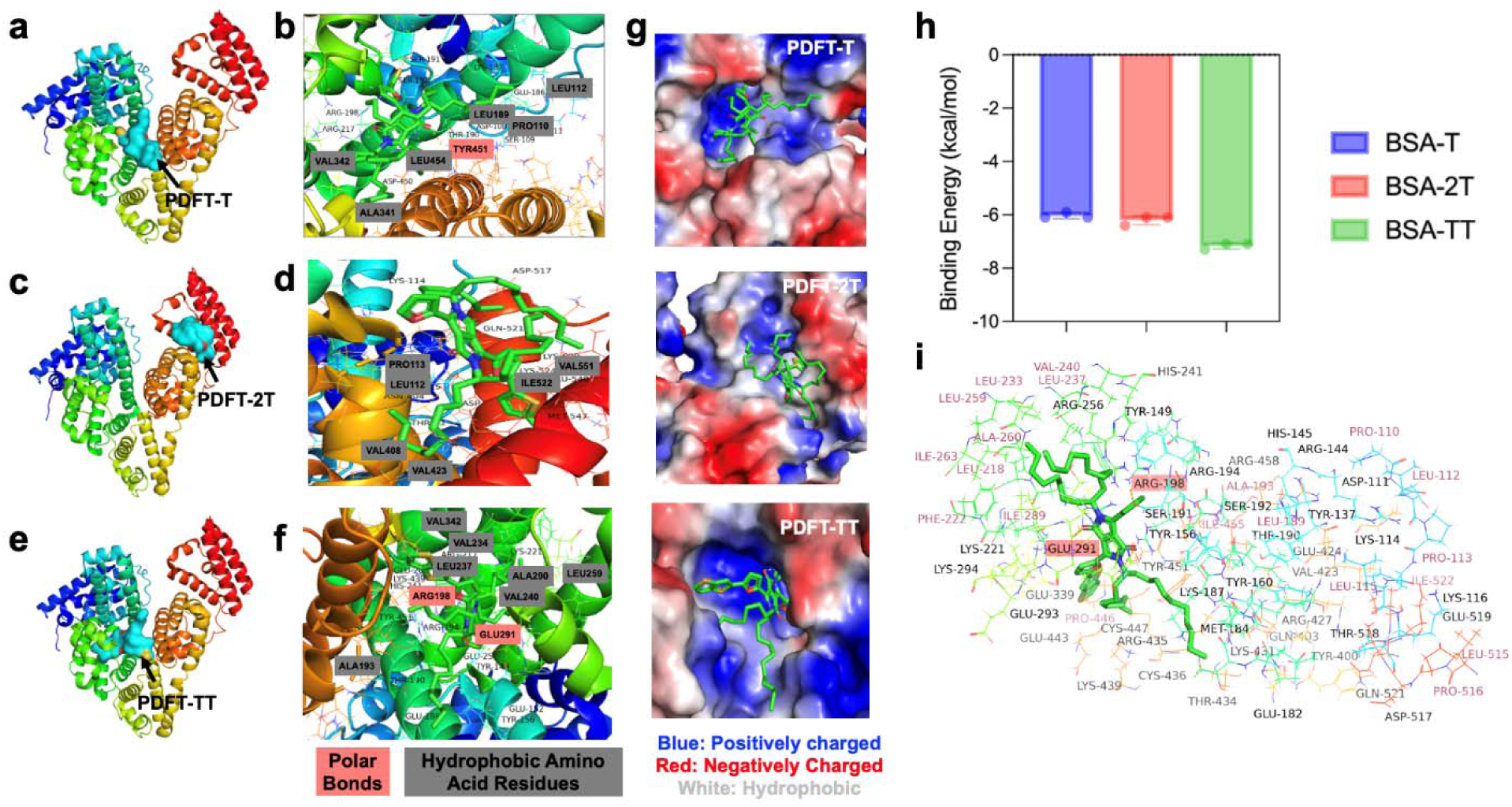
Molecular docking studies to evaluate polymer-protein binding. 3D crystal structure of BSA showing the preferential binding site of (a) PDFT-T, (b) PDFT-2T, and (c) PDFT-TT. The binding interactions and amino acid residues within 4.5 Å of (b) PDFT-T, (d) PDFT-2T, and (f) PDFT-TT. (g) Electrostatic potential mapping of the T-derivative polymer within their respective BSA hydrophobic pocket. The protein pocket is color-coded by electrostatic potential; the potential becomes more positive as the color changes from red to blue. (h) Binding affinity scores (in kcal/mol) with BSA for different T-derivative polymers (n=3). (i) Docking site around the PDFTTT molecule, showing surrounding residues from BSA, depicted as lines. All interacting amino acid residues within the BSA pocket are highlighted (Hydrophilic: Black, Hydrophobic: Red, Polar Bonds: Highlighted in rose red box) surrounding PDFT-TT molecule.

While molecular docking offered useful predictions of possible binding locations and modes of interaction between BSA and PDFT polymers, it’s important to note the inherent limitations of this methodology. Docking generally treats the protein as a rigid structure and does not adequately portray the conformational flexibility of either polymer or BSA, nor the complexity of multivalent binding. Moreover, environmental factors like pH, ionic strength, and temperature can significantly modulate polymer-protein association by stabilizing or destabilizing protein conformations and altering noncovalent interactions. As Bennet et al.^79^ have discussed, such equilibrium binding processes are strongly influenced by the solution environment and can be probed using spectroscopic techniques. It should also be noted that docking was performed with a single repeat unit of the PDFT polymer as a representative ligand, a widely adopted approach in polymer–protein docking studies, since full chain simulations are computationally intractable and unnecessary for capturing local binding interactions. Thus, the docking results presented here are a framework for formulating hypotheses rather than definitive evidence of binding. To complement the predictions from our preliminary docking analysis, we subsequently performed fluorescence titration experiments in a later section, which provide direct experimental evidence of BSA–polymer binding.

### Synthesis, Physicochemical and Photophysical Characterization of BSA-Coated PDFT-Derivative Nanoprobes

For the development of BSA@T/ BSA@TT, or BSA@2T nanoprobes, a modified nanoprecipitation method was used (Figure 4a). PDFT-T/TT/2T polymers were independently dissolved in THF and quickly added dropwise to the BSA containing PBS1X solution under continuous probe sonication. For the development of our control nanoprobes (DSPE-PEG@T, DSPE-PEG@TT, or DSPE-PEG@2T), a routine nanoprecipitation method was adapted (Figure 4a), which involved co-dissolving PDFT-T/TT/2T polymers and DSPE-PEG2k in THF and adding dropwise to PBS1X solution under continuous probe sonication. Once the post-processing was done, they were then subjected to further physicochemical and optical characterizations. The BSA@T, BSA@TT, and BSA@2T nanoprobes exhibited similar hydrodynamic diameters ranging from ∼96 to ∼111 nm, showing no significant differences in the size, polydispersity index (PDI) value, or ζ-potential across polymers (Figure 4b, c, and Figure S8-S10). A comprehensive characterization for all DSPE-PEG@T, DSPE-PEG@TT, and DSPE-PEG@2T nanoprobes was also conducted (Figure S5-S7). Specifically, BSA@TT NPs displayed a hydrodynamic size with an average of 95.44 ± 11 nm and a PDI value of 24.05 ± 4%, while also maintaining their spherical morphology in anhydrous conditions as shown by transmission electron microscopy (TEM) images (Figure 4b, d, and Figure S11). UV-Vis-NIR absorption spectra of BSA@TT nanoprobes were collected, which exhibited a broad absorption center with a major peak at 650 nm (Figure 4e). Subsequently, measuring their NIR-II emission spectra revealed that the NIR-II fluorescence intensity of the BSA@TT increased compared to BSA@2T and BSA@T NPs, showing a similar trend to the fluorescence of the polymers before nanoprobe formation (Figure 2e, 4f). To validate the successful preparation of the BSA-coated nanoprobes, we performed bicinchoninic acid (BCA) protein assay using post-purified BSA@T as a representative formulation. At the characteristic wavelength of 562 nm, BSA@T produced a strong absorbance signal across the well, clearly illustrated by the surface plot and the accompanying purple coloration, confirming the presence of BSA in the nanoprobes. In contrast, DSPE-PEG-coated nanoprobes (DSPE-PEG@T) showed no detectable signal, consistent with the absence of protein (Figure S12). These results demonstrate that BSA was effectively incorporated into the nanoprobe formulation.

**Figure 4.**
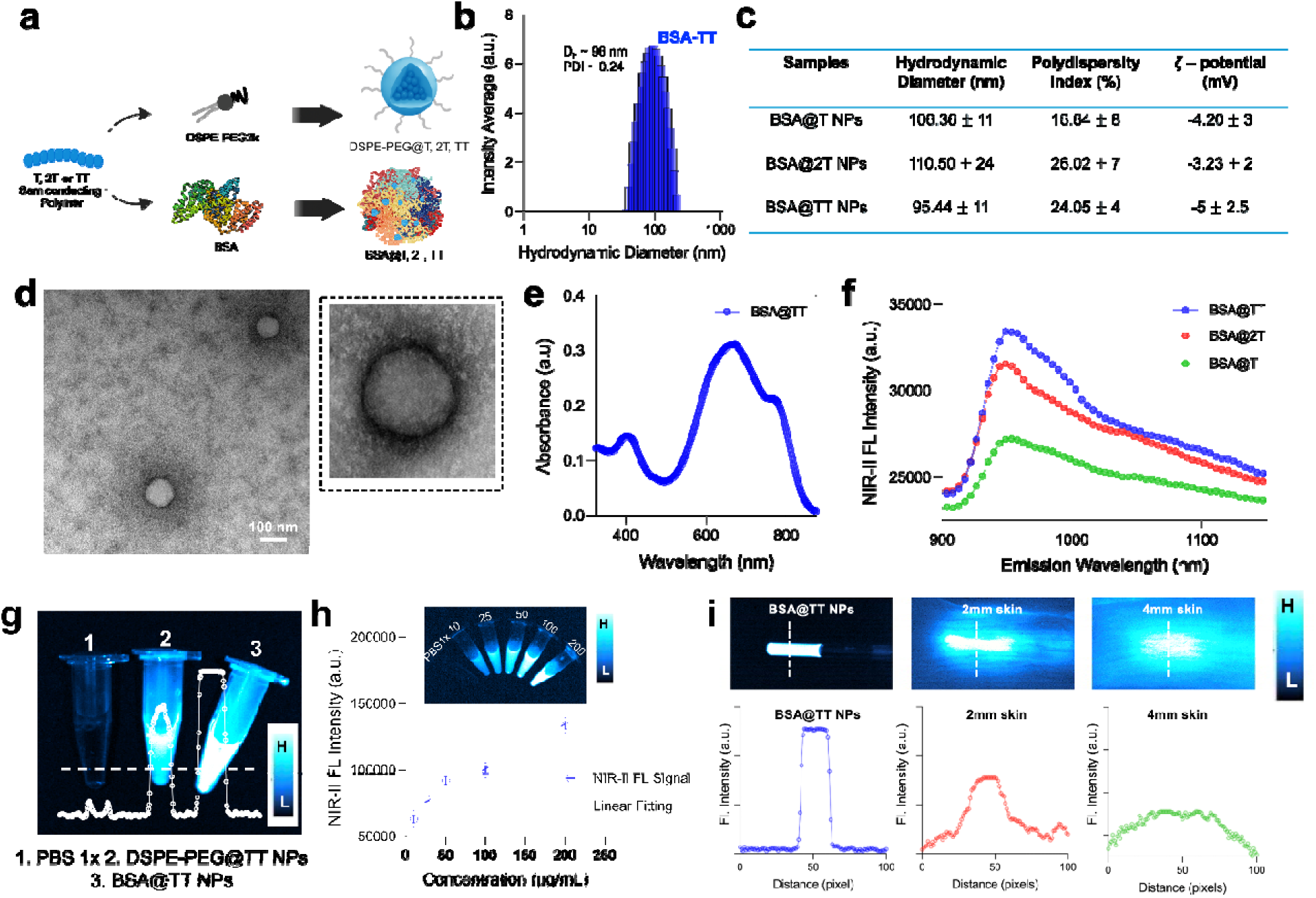
Physicochemical and photophysical characterization of BSA-coated T-derivative NPs. (a) Nanoprecipitation scheme used to synthesize NPs with fluorescent dyes (T, 2T, TT) and stabilizing (DSPE-PEG2k or BSA) amphiphiles. (b) Representative hydrodynamic diameter intensity graph for BSA@TT, measured via dynamic light scattering. (c) Hydrodynamic diameter, PDI, and ζ-potential values for BSA@T, TT, and 2T NPs. (d) Transmission electron microscopy (TEM) images (scale bar: 100 nm) and (e) UV-VIS-NIR absorbance spectra of BSA@TT NPs. (f) NIR-II fluorescence spectra of BSA@T, TT, and 2T NPs (λ_ex_: 808 nm, λ_em_: 900 – 1150 nm). (g) NIR-II fluorescence images with line intensity analysis of PBS1X (control), DSPE-PEG@TT, and BSA@TT (left to right). The dotted line highlighted is used for analysis shown in white. (h) NIR-II fluorescence intensity of BSA@TT NPs at increasing concentrations (10, 25, 50, 100, and 200 μg/mL) with an inset showing the corresponding NIR-II fluorescence image. (i) Tissue penetration analysis with NIR-II fluorescence delineation plots and images of BSA@TT-filled NMR tubes at increasing porcine skin layers (0, 2, and 4 mm). The dotted line highlighted is used for analysis shown in white.

An exhaustive NIR-II emission profile for all polymers and their lipid and protein NP formulations was also collected for excitation wavelengths 397, 660, 709, and 790 nm (Figure S13). Excitation-dependent PL spectra further highlight how encapsulation modulates the optical properties of the polymers. Compared to TT in THF, BSA@TT exhibited enhanced long-wavelength emission (>1100 nm) (Figure S13), consistent with reduced aggregation-induced quenching and stabilization of the polymer chains within the albumin matrix.^52^ DSPE-PEG assemblies, by contrast, showed broader and weaker emission. These findings underscore how the choice of encapsulation strategy, particularly protein nanoengineering, directly tunes emission profiles and enhances brightness for deep-tissue imaging. To further illustrate the amplified NIR-II fluorescence performance of the PDFT-TT polymer when coated with either a lipid (DSPE-PEG) or a functional protein (BSA), NIR-II fluorescence imaging was conducted using the IR Vivo Imager. Notably, BSA@TT (200 µg/mL) exhibited significantly stronger NIR-II brightness, outperforming DSPE-PEG@TT (200 µg/mL), showcasing the unique advantage of using a functional albumin matrix (Figure 4g).

Moreover, a strong positive correlation was observed between NIR-II fluorescent intensity and BSA@TT nanoprobe concentration, which was systematically varied from 200 µg/mL to 10 µg/mL, yielding an R^2^ value of 0.98 (Figure 4h). Fluorescence titration experiments provided quantitative support for the molecular docking predictions of BSA-TT interactions. The binding constant (K_d_ = 75.96 µM) lies within the expected range of albumin-ligand interactions and reflects a favorable affinity that is sufficiently strong to ensure stable incorporation of TT into the BSA matrix. The high maximum binding capacity (*B*_max_ = 6051 a.u.) further underscores albumin’s ability to accommodate multiple TT molecules, consistent with its well-established role as a versatile carrier of hydrophobic ligands (Figure S14). The encapsulation efficiency of BSA@TT determined by UV–Vis calibration was 58 ± 5%, confirming effective incorporation of TT into the albumin nanoprobes (Figure S15). Continuous irradiation studies further confirmed that BSA@TT retained stable fluorescence over 2 h of laser exposure, whereas ICG rapidly photobleached, underscoring the superior photostability of the albumin-engineered probes. (Figure S16) Together, these findings confirm BSA as an effective encapsulation matrix for TT and provide a mechanistic rationale for favorable stability and potential optical enhancement for BSA@TT nanoprobes.

To further assess the bioimaging potential of BSA@TT’s, NIR-II fluorescent intensities were measured through increasing tissue depths. Impressively, BSA@TT (200 µg/mL) generated a detectable signal through more than 4mm of porcine skin and resolved the contours of the NMR tube containing the solution (Figure 4i). Our data further demonstrate that BSA@TT nanoprobes retain detectable emission through porcine muscle sections up to 8 mm thick, underscoring their relevance for intraoperative deep-tissue imaging. (Figure S17) The ability of BSA@TT nanoprobes to maintain a strong, localized fluorescence signal beneath the tissues highlights their resistance to the light scattering and absorption typically encountered in biological environments. The BSA@TT nanoprobes also demonstrated robust colloidal stability under conditions that mimic physiological and pathological environments. Dynamic light scattering measurements showed that the hydrodynamic diameter remained essentially unchanged when incubated for one week in buffers of varying pH (4.5, 5.5, and 6.5, representing endosomal, tumor, and mildly acidic microenvironments, respectively), at 37 °C, and in the presence of plasma (Figure S18). Additionally, BSA@TT nanoprobes retained their fluorescence intensity upon incubation with FBS, plasma, or porcine blood, with no statistically significant differences compared to PBS (Figure S19). These results confirm that BSA@TT maintains its structural integrity and dispersion stability under physiologically relevant conditions, supporting its suitability for use in living tissues. Notably, their NIR-II optical properties remained largely unaffected upon interaction with porcine blood, a medium known to quench the fluorescence of many investigational nanoprobes, likely due to several factors: (i) robust surface passivation offered by BSA, (ii) their emission beyond the primary absorption range of hemoglobin, and (iii) inherent resistance to aggregation and biological quenchers.^80^

The optical properties of DSPE-PEG@TT, and BSA@TT nanoprobes were evaluated, with key parameters including quantum yield (</), mass extinction coefficient ({_m_), and brightness (B) summarized in Table S1. ICG, a clinically approved dye, has its modest quantum yield (1.7%).^60^ Both DSPE-PEG@TT and BSA@TT nanoprobes exhibited significantly amplified optical performance, despite their SP content being tested at much lower concentrations (5.54 × 10^-7^ mol/L). DSPE-PEG@TT displayed a mass extinction coefficient of 15,230 L/mg.cm, resulting in a brightness of 418.83. BSA@TT outperformed DSPE-PEG@TT with a similar extinction coefficient of 15,230 L/mg.cm, leading to an even greater brightness of 581.79, which is ∼1.5 times brighter than DSPE-PEG@TT (Table S1). The fluorescent quantum yield (φ) values of the isolate PDFT-T, PDFT-TT, and PDFT-2T polymers are 0.16%, 0.40%, and 0.30%, respectively, in reference to ICG (Figure S21). Notably, BSA encapsulation enhanced the quantum yield of TT to 3.82%, compared to 2.75% for DSPE-PEG@TT and 0.40% for TT in THF (Figure S20-S21), underscoring the advantage of protein nanoengineering in boosting brightness and stability.

The mass extinction coefficient (ε_m_) describes the absorbance of a substance per unit mass at a given wavelength. While this parameter reflects how strongly a sample absorbs light, fluorescence brightness is determined by the product of ε_m_ and the quantum yield (</), which quantifies the efficiency with which absorbed photons are converted into emitted fluorescence. Consequently, samples with comparable mass extinction coefficients can exhibit markedly different fluorescence intensities if their quantum yields differ. In the case of BSA@TT displaying greater fluorescence brightness than DSPE-PEG@TT despite similar ε_m_ values, this is likely due to the BSA matrix providing a more favorable microenvironment for the polymer. This stabilization may reduce non-radiative decay pathways and enhance quantum yield.^44,66^ Additionally, BSA may mitigate aggregation or quenching effects that can occur in DSPE-PEG@TT systems. These results demonstrate the importance of the nanoprobe design and the surrounding chemical environment in tuning the optical properties of NIR-II fluorescent nanoprobes. BSA encapsulation provides a promising strategy to maximize fluorescence brightness for improved bioimaging applications.

### Tumor-Margin Delineation in *Ex Vivo* Porcine Tissues Embedded with Tumor-Mimicking Phantoms Containing BSA@TT Nanoprobes

To accurately assess the bioimaging potential of BSA@TT nanoprobes under conditions that mimic image-guided surgery, it is critical to replicate the complex optical properties of biological tissues. In real surgical scenarios, imaging agents must perform within an environment that includes endogenous fluorophores, high scattering, and strong absorption – all of which interfere with fluorescence detection. These challenges can be addressed by shifting the fluorescence emission of our imaging probes further into the NIR-II (1000 – 1600 nm) range, where minimal spectral overlap is observed from such components (endogenous fluorophores, scattering, absorption). In this study, we considered the complex optical environment that our nanoprobe would encounter once at a tumor-tissue site by developing tumor-mimicking phantoms and incorporating intralipid (light-scattering component) and hemoglobin (light-absorbing component) with BSA@TT nanoprobes (NIR-II fluorescence component) (Figure 5a). This strategy, previously reported by our group,^61, 62^ offers a reproducible, cost-effective, and adaptable platform for evaluating the optical performance of a myriad of dyes across a range of surgical applications.

**Figure 5.**
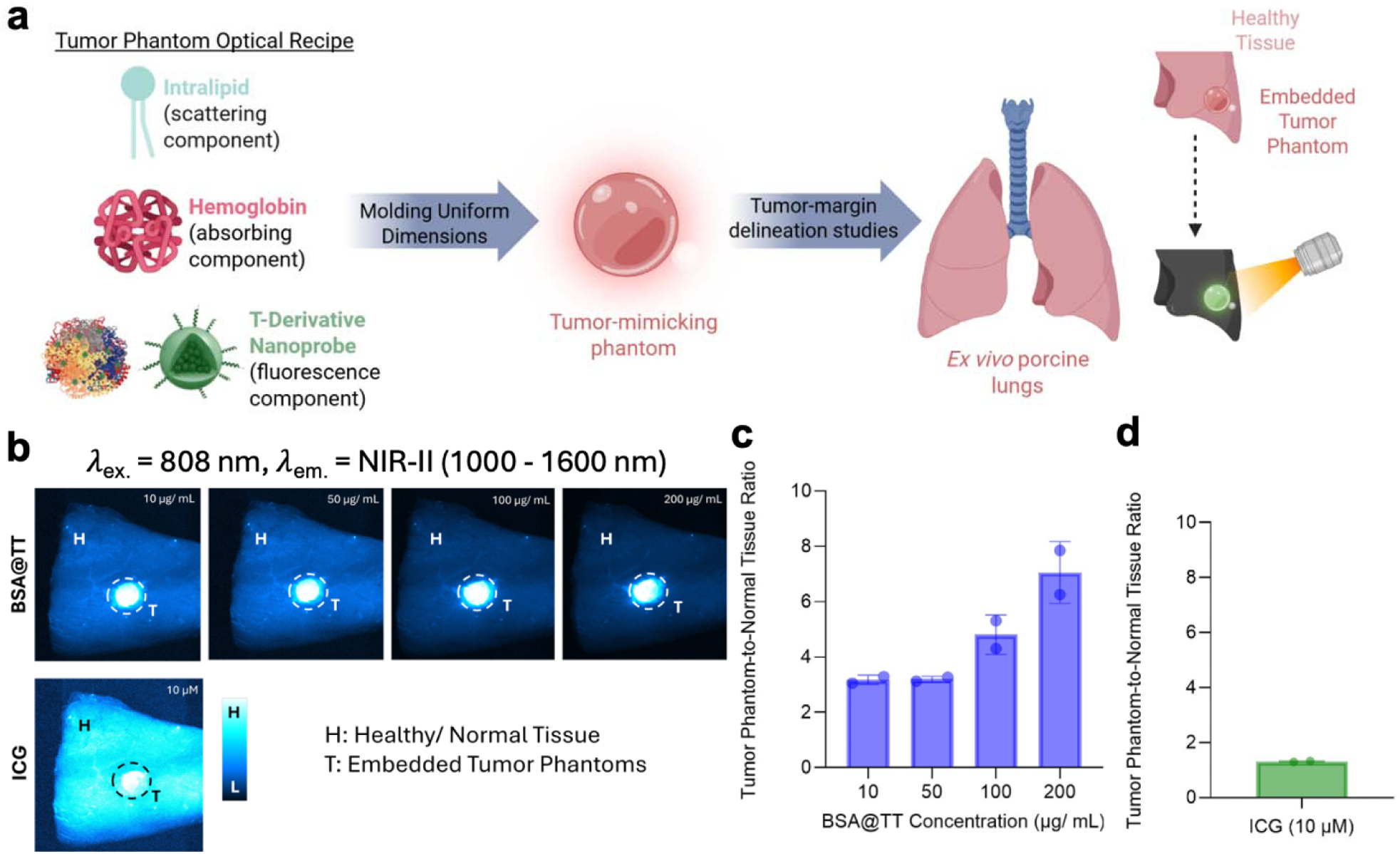
Image-guided surgery simulation using *ex vivo* porcine lung tissue with embedded tumor-mimicking phantoms. (a) Schematic illustration depicting tumor phantom synthesis and its use in tissue discrimination studies. (b) NIR-II fluorescence imaging of embedded tumor phantoms incorporating BSA@TT at increasing concentrations (10, 50, 100, and 200 µg/mL) and ICG (10 µM) with their respective tumor-to-normal tissue ratios (c, d).

For anatomical context, tumor-mimicking phantoms were embedded within freeze-dried porcine lung tissue to simulate buried solid tumors surrounded by healthy parenchyma, a common scenario for solid tumor cases.^81^ To evaluate the ability of BSA@TT NP-containing tumor phantoms to differentiate from surrounding healthy tissues, we incorporated BSA@TT nanoprobes at increasing concentrations (10, 50, 100, and 200 µg/mL) within a tumor-mimicking phantom (Figure 5b). To establish a control, a tumor-mimicking phantom was developed comprising the usual elements and indocyanine green (ICG). Our workflow used DSPE-PEG@TT as a screening control to select the superior encapsulation matrix (BSA) and then benchmarked BSA@TT against ICG, the clinical standard, in large-organ ex vivo models to emphasize translational relevance. Although ICG is an FDA-approved fluorescent dye and is commonly used for pre-clinical tumor surgeries, it is susceptible to a detrimental amount of photobleaching and spectral interference in complex tissue environments. All BSA@TT concentrations exhibited tumor phantom-to-normal tissue fluorescence ratios of at least 3.81 (n = 2), significantly outperforming ICG (10 µM) in the NIR-II window, which exhibited a tumor phantom-to-normal tissue ratio of 1.34 ±0.06 (n=2) (Figure 5c, d). Specifically, at 10 µg/mL and 200 µg/mL, BSA@TT exhibited 2.9-fold and 6.6-fold higher contrast ratios than ICG, respectively. All four BSA@TT concentrations similarly outperformed ICG (10 µM) in terms of tumor phantom-to-normal tissue fluorescence ratios within the NIR-I and LP1250 emission windows (Figure S22). These results provide quantitative evidence that BSA@TT nanoprobes enable more accurate delineation of the tumor boundaries under optically realistic conditions, highlighting their potential as a next-generation imaging agent for solid tumor visualization.

### Comparative Study of Microvasculature Visualization in *Ex Vivo* Ovine Brain using BSA@TT Nanoprobes

Building upon our initial efforts to simulate buried tumor lesions, we further investigated the potential of BSA@TT nanoprobes to delineate anatomical features in *ex vivo* ovine brain models. Given that ICG is widely used in clinical settings for visualizing organ-specific blood flow and assessing tissue perfusion, we sought to compare its imaging performance directly with BSA@TT nanoprobes under similar experimental conditions. We first injected *ex vivo* ovine brains with 100 µL of both ICG (10 µM, clinical dose)^82^ and BSA@TT (200 µg/mL) (Figure 6a). Notably, BSA@TT demonstrated superior performance over ICG across all three NIR-II emission windows (925-1000, >1000, and > 1250 nm) in microvasculature structures. Line profile analysis of fluorescence intensity across the injection sites revealed distinct intensity peaks that appeared for the BSA@TT nanoprobes, showcasing their capability to resolve microvasculature networks extending beyond the sulci (indented brain fissures). Upon excitation with an 808 nm laser, BSA@TT nanoprobes successfully identified four discrete anatomical structures within the 925-1000 and > 1250 nm emission windows and two structures within the >1000 nm emission window (Figure 6b-e). In contrast, under identical imaging conditions and the same brain model (to maintain consistent vasculature structure and structure density), ICG exhibited markedly weaker performance, with lower signal-to-background and limited visualization compared to BSA@TT nanoprobes. It is well established that ICG fluorescence can increase upon binding to albumin in vivo; however, despite this effect, ICG remains constrained by aggregation-induced quenching, rapid clearance, and poor photostability performances under imaging conditions. By contrast, the BSA@TT nanoprobes integrate albumin intrinsically, yielding enhanced brightness together with improved stability and imaging performance.

**Figure 6.**
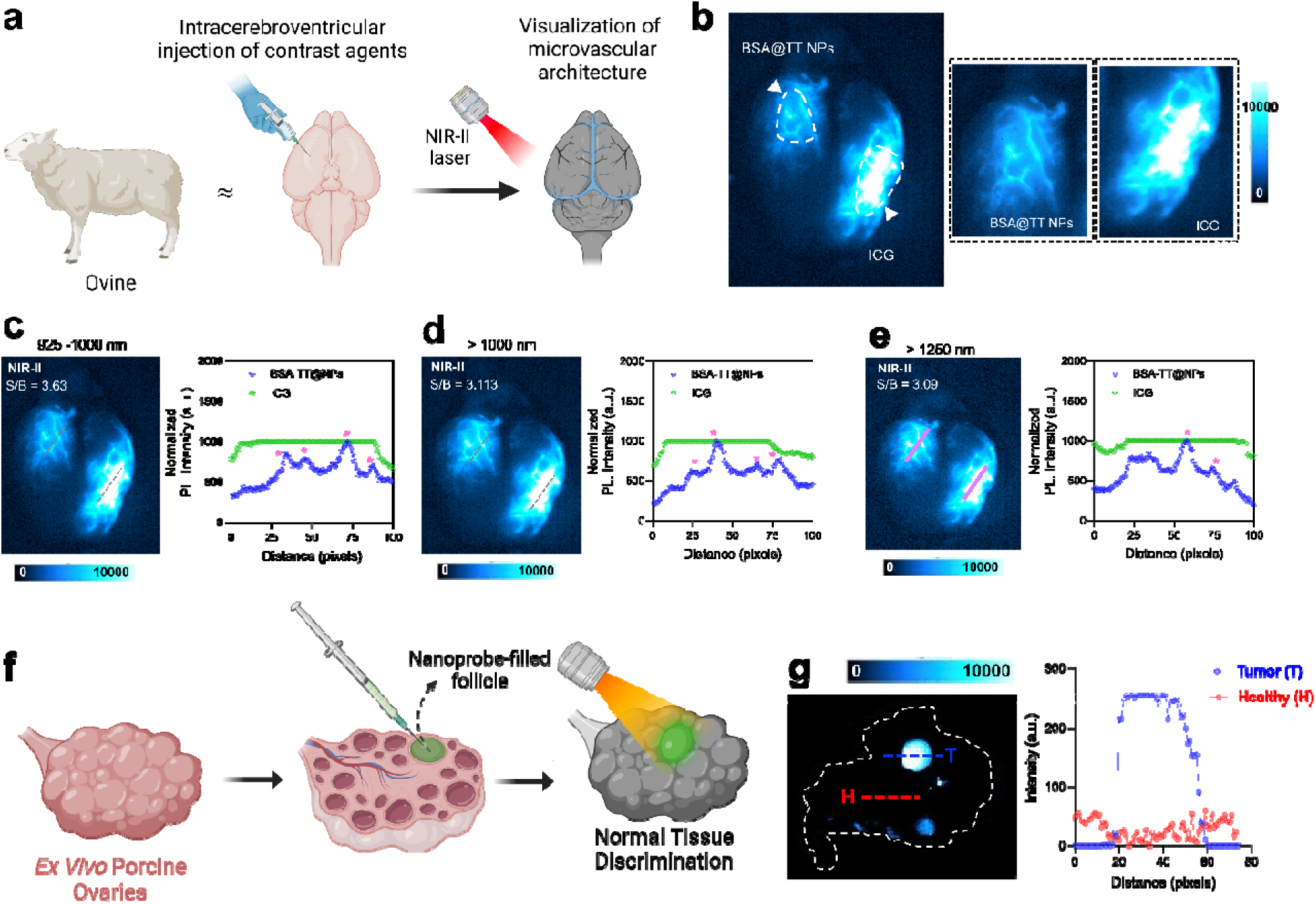
Microvasculature visualization and tissue discrimination studies using *ex vivo* ovine brain and porcine ovaries. (a) Schematic illustration of the *ex vivo* ovine brain injection method used for microvasculature visualization. (b) Representative NIR-II fluorescence images of location-specific injections of BSA@TT NP solution and ICG dye, at *in vivo* -representative concentrations. *Ex vivo* ovine brain NIR-II fluorescence images (λ_ex_: 808 nm) and microvasculature delineation plots of BSA@TT and ICG injections with emission windows (c) 925 – 1000 nm, (d) >1000 nm, and (e) >1250 nm, n=3. (f) Schematic illustration of the *ex vivo* porcine ovary injection method used for tissue discrimination analysis. (g) Representative NIR-II fluorescence image and intensity graph of BSA@TT-filled porcine ovary follicle, n = 3 injections (λ_ex_: 808 nm, λ_em_: 925-1000 nm).

While BSA@TT resolves discrete microvascular features ex vivo and yields SBRs of 3.63, 3.11, and 3.09 (925–1000, >1000, >1250 nm, respectively), these measurements are obtained in non-perfused tissue and therefore do not capture hemodynamic effects (flow, wash-in/wash-out, extravasation) that influence in vivo contrast dynamics. Accordingly, we have replaced qualitative color bars with quantitative intensity scales and annotated SBR values to convey dynamics more transparently. Future work will include PK and biodistribution (t½, AUC, clearance route) to substantiate translation as a contrast agent; however, that is outside the scope of the current work.

### Discrimination Between Tumor and Healthy Tissue in *Ex Vivo* Porcine Ovaries *via* BSA@TT Nanoparticles

To further investigate the extent of our BSA@TT’s nanoprobes tissue discrimination abilities, we conducted a comparative imaging study using *ex vivo* porcine ovaries as a tumor-mimicking model. *In vivo,* nanoprobe accumulation in ovarian tumors can occur due to enhanced vascular permeability, irregular angiogenesis, and impaired lymphatic drainage which are the hallmarks of the enhanced permeability and retention (EPR) effect commonly leveraged by nanoprobe-based cancer bioimaging.^83–85^ To simulate this localized accumulation in a controlled setting, 100 µL of BSA@TT (200 µg/mL) and ICG (10 µM) was directly injected into the follicular cavity of a mature ovary follicle (Figure 6f). The antral space in these follicles provides an ideal structurally relevant reservoir, while the ∼3mm of overlying tissue offers a realistic depth for assessing NIR-II signal penetration and resolution.^86^ For all line profiles (Figures S23–S24), dashed ROIs were drawn across tumor (T) and adjacent healthy (H) regions to systematically compare contrast rather than randomly selected lines. This comparative study revealed that BSA@TT nanoprobes precisely delineated the injection site and produced fluorescent intensities 4.25 times higher than the surrounding ovarian tissue within the 925 – 1000 nm emission window (Figure 6g). Additional NIR-II imaging of this same ovary revealed fluorescent intensities 2.09 and 3.04 times greater than surrounding tissues within the emissions windows >1000 and >1250 nm, respectively (Figure S23). Conversely, ICG injections resulted in poor spatial localization, low contrast, and inconsistent fluorescence profiles across all emission windows (Figure S24). These experiments further showcase that BSA@TT nanoprobes provide significantly improved anatomical contrast and tissue discrimination compared to FDA-approved ICG, supporting their potential for tumor localization and margin assessment in cancer bioimaging.

### *In Vitro* Cytotoxicity Assessments and Cellular Internalization of BSA@TT Nanoprobes

Upon a rigorous physicochemical and NIR-II optical characterization of BSA@TT nanoprobes and their counterpart DSPE-PEG@TT nanoprobes, we next attempted to assess their biocompatibility and ability to be internalized by *in vitro* OVCAR8 ovarian tumor cells. Both DSPE-PEG@TT and BSA@TT nanoprobes exhibited minimal toxicity in the OVCAR8 cells, even at high concentrations of 100 µg/mL (Figure S25). To examine their cellular internalization, we utilized fluorescence spectroscopy and IVIS imaging to quantify uptake into OVCAR8 cells. Both DSPE-PEG@TT and BSA@TT nanoprobes demonstrated a time-dependent cellular uptake with the maximum uptake at 12 hours for both the probes, but BSA@TT nanoprobes showed significantly higher uptake compared to DSPE-PEG@TT nanoprobes (Figure S26). Such a trend could be attributed to the presence of secreted proteins acidic and rich in cysteine (SPARC) receptors that are highly expressed on metastatic tumor cells, including OVCAR8, which are known to bind to albumin.^87–89^ These receptors likely facilitate a selective endocytosis uptake mechanism for BSA@TT nanoprobes. Preliminary *in vitro* assays confirmed biocompatibility and cellular uptake of BSA@TT in ovarian adenocarcinoma cells (Figure S26), supporting their suitability for *ex vivo* bioimaging applications.

### Hemocompatibility Assessments of BSA@TT Nanoprobes

Evaluating the interactions between nanoprobes and red blood cells is an important step in the preliminary assessment of biocompatibility. To this end, we quantified the level of hemolysis, defined by the erythrocyte cell destruction, that occurred when porcine red blood cells were incubated with our BSA@TT NPs at increasing concentrations (1, 5, 10, 50, and 100 µg/mL) (Figure 7a). Hemolysis assays were conducted in parallel using standard controls: phosphate-buffered saline (PBS1X) as a negative control, and deionized water as a positive control (Figure 7b). All BSA@TT nanoprobes-treated samples retained red blood cell pellets at the bottom of the microcentrifuge tube, indicating the preservation of cell integrity. In contrast, the DI water control resulted in complete hemolysis, evidenced by a uniformly red supernatant and the absence of a cell pellet (Figure 7c). PBS-treated cells formed a dense pellet with a clear supernatant, consistent with minimal hemolysis. The BSA@TT nanoprobes treated samples of all concentrations induced hemolysis levels below the 10% threshold for blood biocompatibility, with the lowest rate observed at 10 µg/mL (3.12 +/-1.08%) (Figure 7d). These results confirm that BSA@TT nanoprobes are hemocompatible across the tested dose range and unlikely to induce red blood cell lysis under physiological conditions.

**Figure 7.**
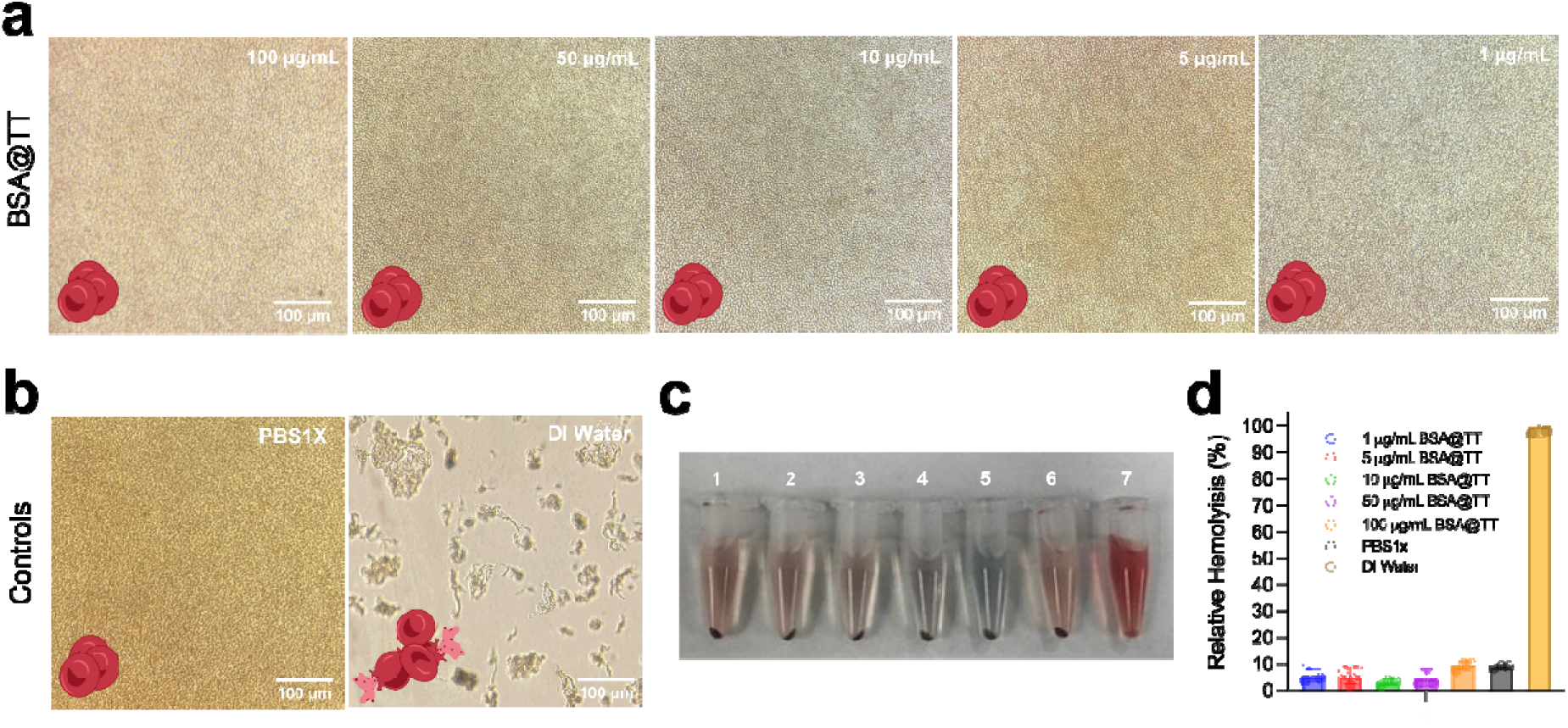
Hemocompatibility studies of BSA@TT NPs with porcine blood. Inverted microscopic images at 200X magnification of a) BSA@TT NPs (100, 50, 10, 5, and 1 µg/mL), b) PBS1X (negative control), and DI water (positive control) after one hour of incubation at 37°C with porcine blood. c) White light image of RBC pellets from each incubated sample: (1) 100 µg/mL, (2) 50 µg/mL, (3) 10 µg/mL, (4) 5 µg/mL, (5) 1 µg/mL, (6) PBS1X, and (7) DI water. (d) Relative hemolysis percentages of all BSA@TT treatment and control groups.

These preliminary findings showcase negligible *in vivo* toxicity to healthy porcine blood, at least within our tested dosage range, making them suitable for potential *in vivo* applications. However, further investigations are warranted to systematically explore both short-term and long-term toxicity of these nanoprobes across various doses in *in vivo* models, but that is beyond the scope of the current study.^90, 91^

## CONCLUSIONS

In summary, we have developed a next-generation NIR-II imaging nanoprobe platform through two complementary techniques: molecular engineering and protein nanoengineering. By optimizing the placement of thiophene units within the semiconducting polymer backbone and leveraging bovine serum albumin as a biologically functional encapsulation matrix, we generated ultrabright, colloidal stable nanoprobes with superior imaging performance compared to the FDA-approved dye, ICG. These BSA@TT nanoprobes exhibited amplified NIR-II fluorescence brightness, excellent colloidal stability in biologically relevant environments, and deep tissue penetration, significantly outperforming conventional lipid-coated nanoprobes (DSPE-PEG@TT) or ICG. Their imaging capability was validated in *ex vivo* large-animal models across diverse applications, including buried tumor lesion localization, tumor margin delineation, and microvascular visualization, demonstrating their strong potential for clinical translation in image-guided surgery and diagnostics. This dual-engineering approach offers a robust and versatile framework for advancing NIR-II nanoprobe development towards real-world bioimaging applications.

## ASSOCIATED CONTENT

### Supporting Information

A listing of the contents of each file supplied as Supporting Information should be included. For instructions on what should be included in the Supporting Information as well as how to prepare this material for publications, refer to the journal’s Instructions for Authors. The following files are available free of charge.

### Author Contributions

I.S. conceived the idea and designed all the experiments. I.V. led the study and was involved in data collection and curation, formal analysis, investigation, methodology, validation, and visualization of all the experiments under the supervision of I.S. Synthesis and characterization of T, TT, and 2T was conducted R.P. under the supervision of J.T. NIR-II fluorescence spectra, and QY were collected by N.G. under the supervision of J.T. Processing and physicochemical characterization of DSPE-PEG@T, TT, 2T, and BSA@T, TT, 2T were performed by I.V., and R.R. under the supervision of I.S. Molecular docking studies were conducted by A.H. *Ex vivo* tissue studies were performed by I.V. and A. H. under the supervision of I.S. and U.B. *In vitro* toxicity studies and cellular uptake were performed by I.V. under the supervision of I.S. Hemocompatibility studies were conducted by R. R. and I.V. under the supervision of I.S. The manuscript was written by I.V. and I.S. with edits from J.T. All authors have approved the final version of the manuscript.

## ACKNOWLEDGMENTS.

I.S. acknowledges the Edward E. Whitacre Jr. College of Engineering and Texas Tech University for research support. Imaging studies on the IR VIVO system were funded by the Core Facility Support Award (grant number RP200572) from the Cancer Prevention and Research Institute of Texas (CPRIT) to the Imaging Core, Texas Tech University Health Sciences Center at Amarillo. The authors acknowledge Md. Hasnat Rashid for acquiring TEM images. All graphics in this paper were made in BioRender.

