## Supplemental File for "Ultrabright NIR-II Nanoprobes for *Ex Vivo* Bioimaging: Protein Nanoengineering Meets Molecular Engineering"

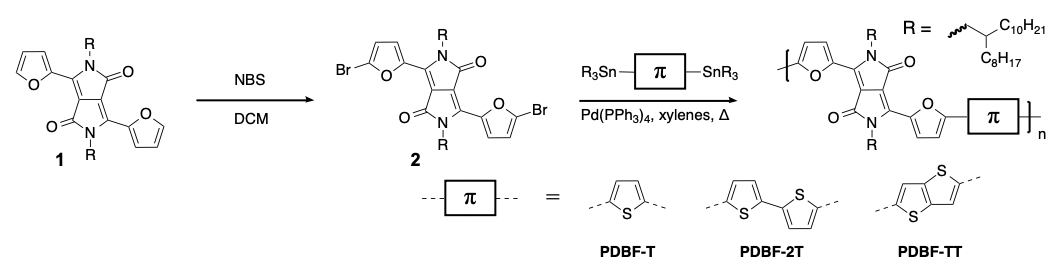

**Scheme S1**. Monomer synthesis and polymerization of PDFT-T, PDFT-2T, and PDFT-TT copolymers with branched aliphatic side chains via Stille polycondensation.

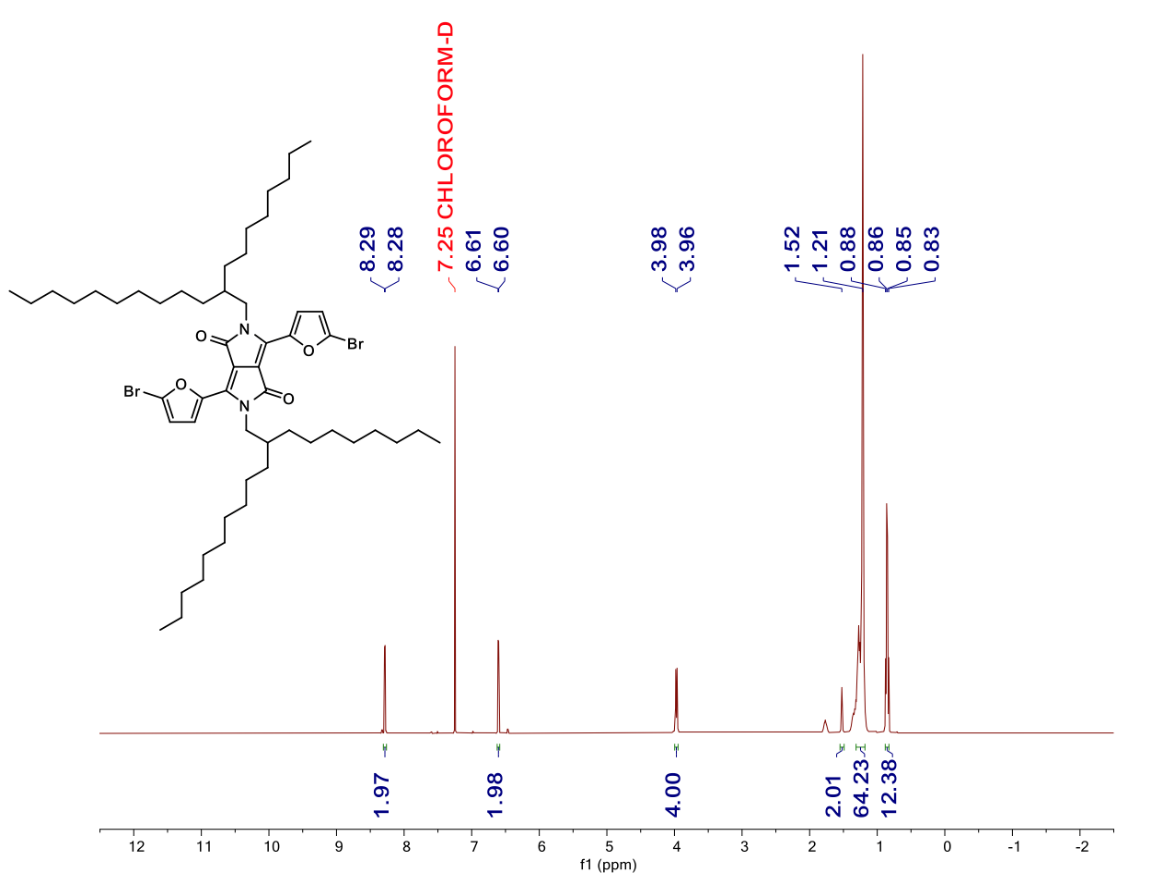

**Figure S1.** ^1^H NMR spectrum of 3,6-bis(5-bromofuran-2-yl)-2,5-bis(2-octyldodecyl)-2,5-dihydropyrrolo[3,4-c]pyrrole-1,4-dione (**2**).

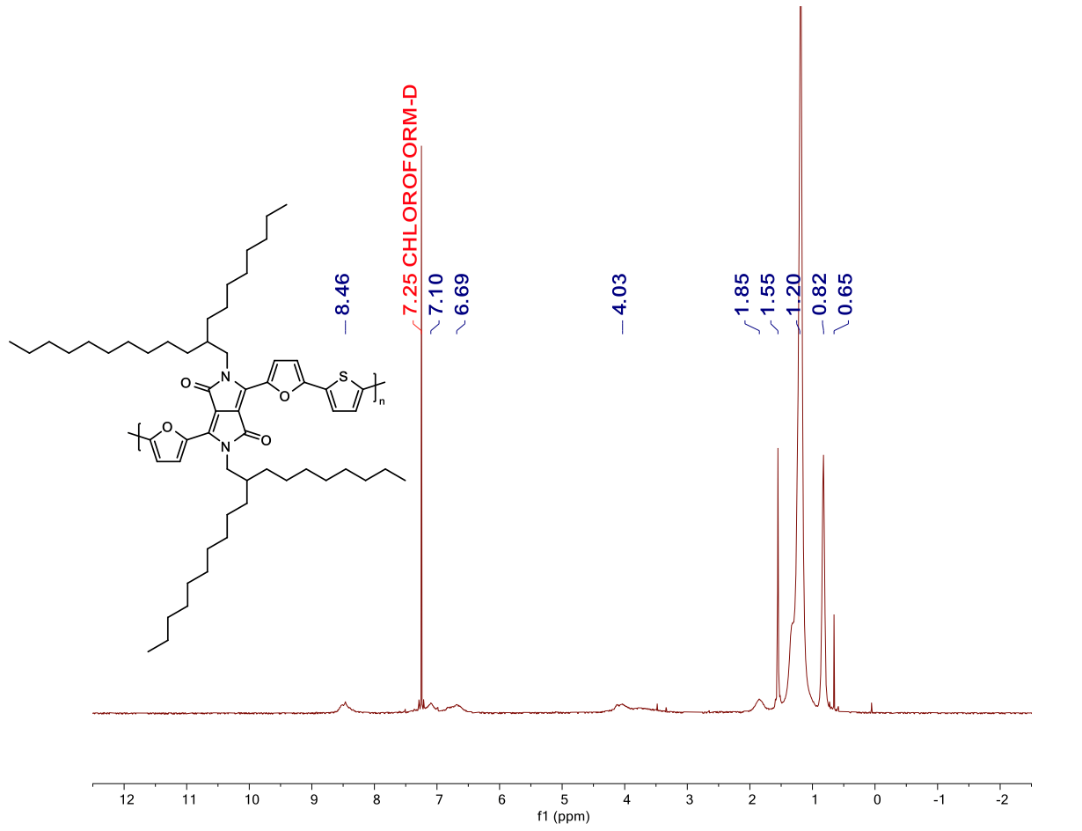

**Figure S2.** ^1^H NMR spectrum of PDFT-T.

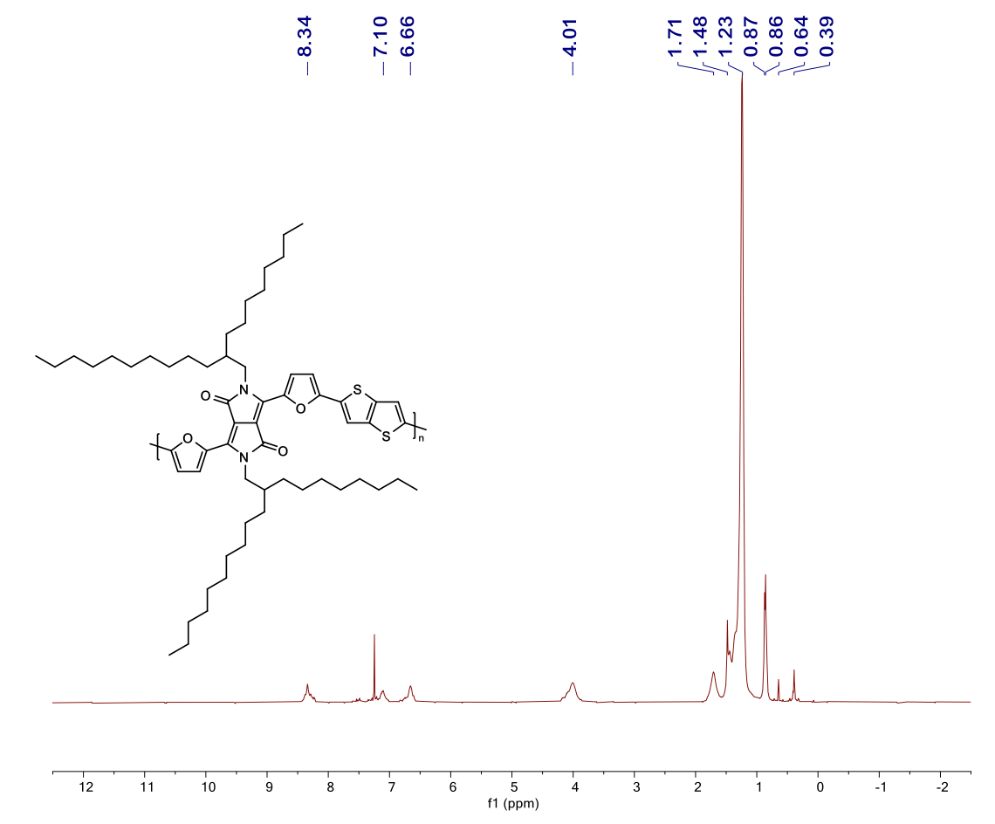

**Figure S3.** ^1^H NMR spectrum of PDFT-TT.

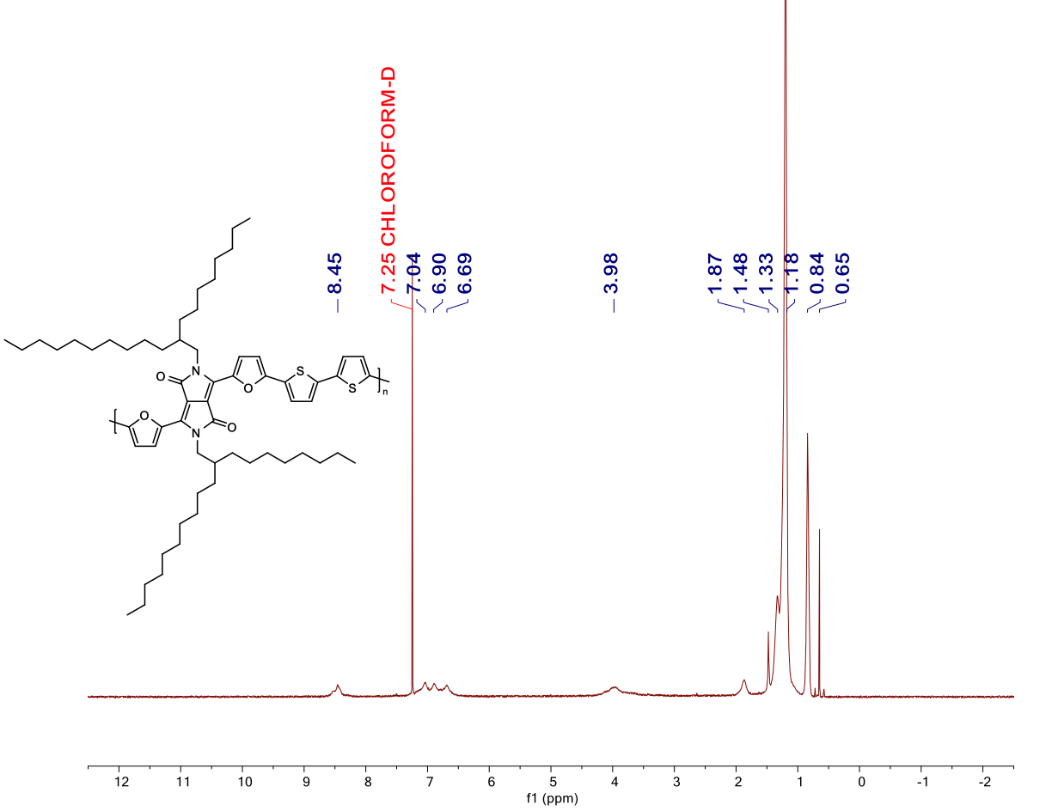

**Figure S4.** ^1^H NMR spectrum of PDFT-2T.

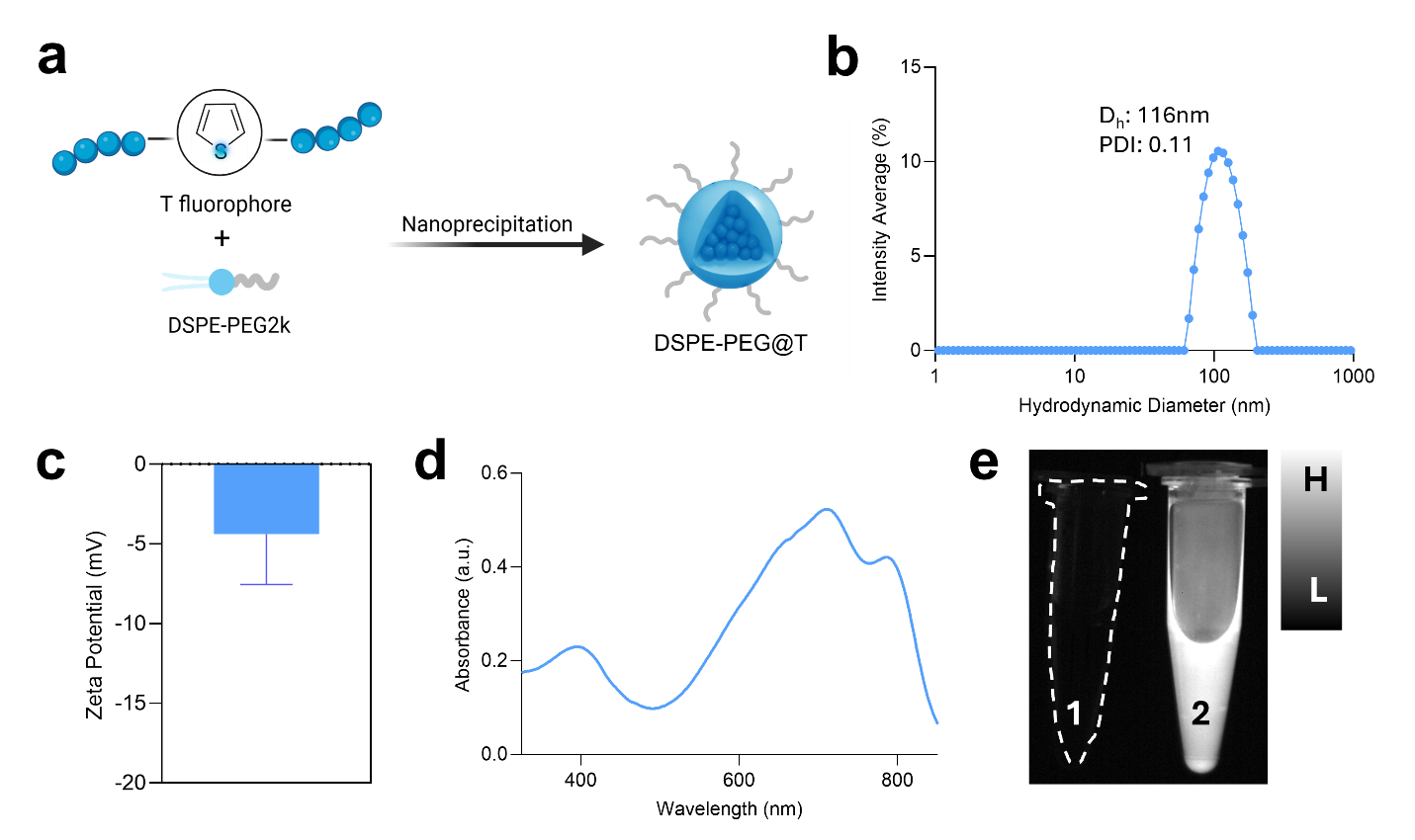

**Figure S5. Physiochemical characterization of DSPE-PEG@T nanoparticles.** (a) Schematic representation of the preparation of DSPE-PEG@T. (b) Hydrodynamic diameter and PDI measurements *via* dynamic light scattering. (c) Zeta potential, in mV. (d) UV-Vis-NIR absorbance spectra. (e) IR Vivo fluorescence image of PBS1X (1) and DSPE-PEG@T nanoparticles (2) with a $\lambda$_ex._ = 808 nm and $\lambda$_em._ = 1000 – 1600 nm. Corresponding scale bar is provided for comparison.

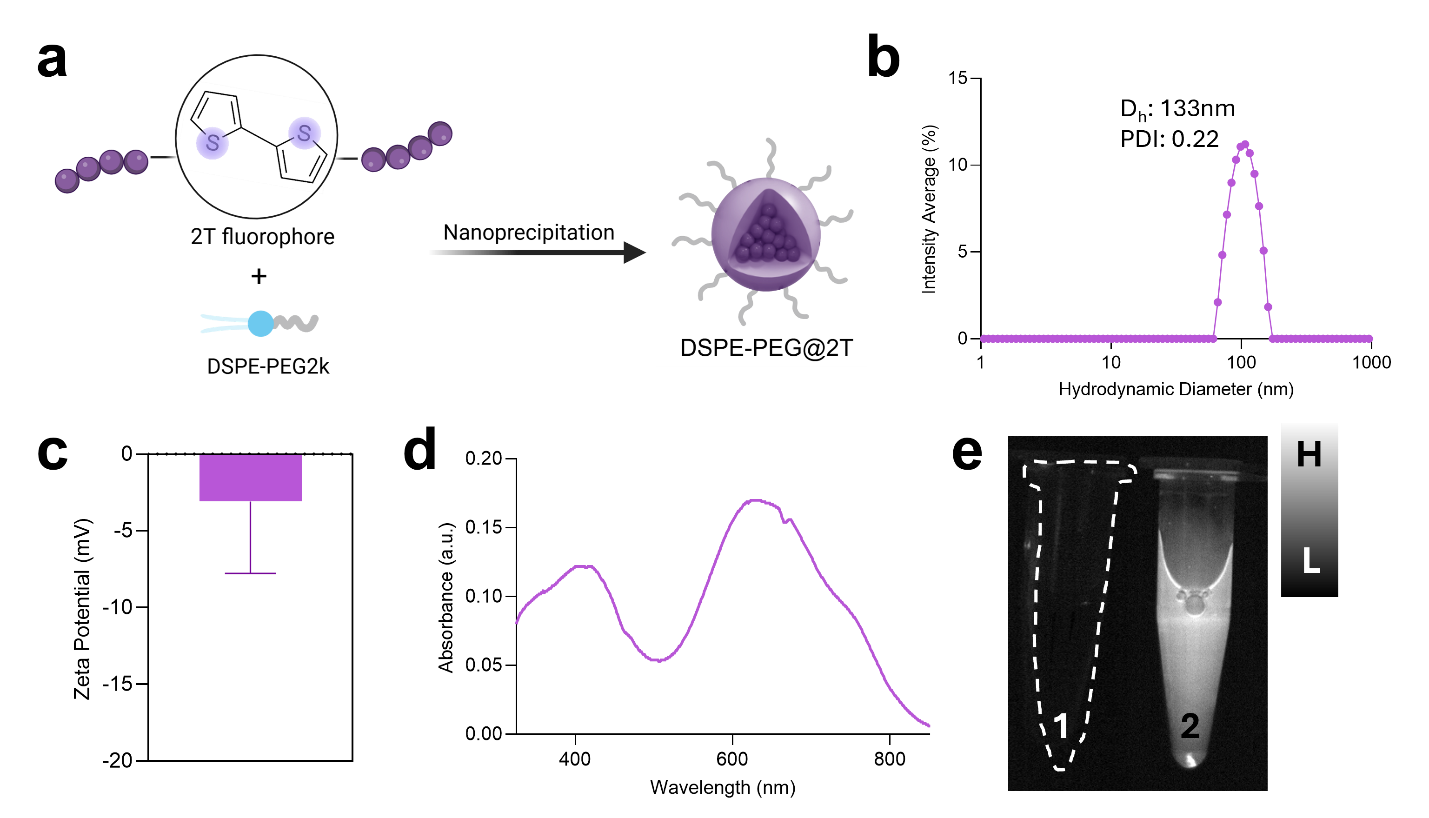

**Figure S6. Physiochemical characterization of DSPE-PEG@2T nanoparticles.** (a) Schematic representation of the preparation of DSPE-PEG@2T. (b) Hydrodynamic diameter and PDI measurements *via* dynamic light scattering. (c) Zeta potential, in mV. (d) UV-Vis-NIR absorbance spectra. (e) IR Vivo fluorescence image of PBS1X (1) and DSPE-PEG@2T nanoparticles (2) with a $\lambda$_ex._ = 808 nm and $\lambda$_em._ = 1000 – 1600 nm. Corresponding scale bar is provided for comparison.

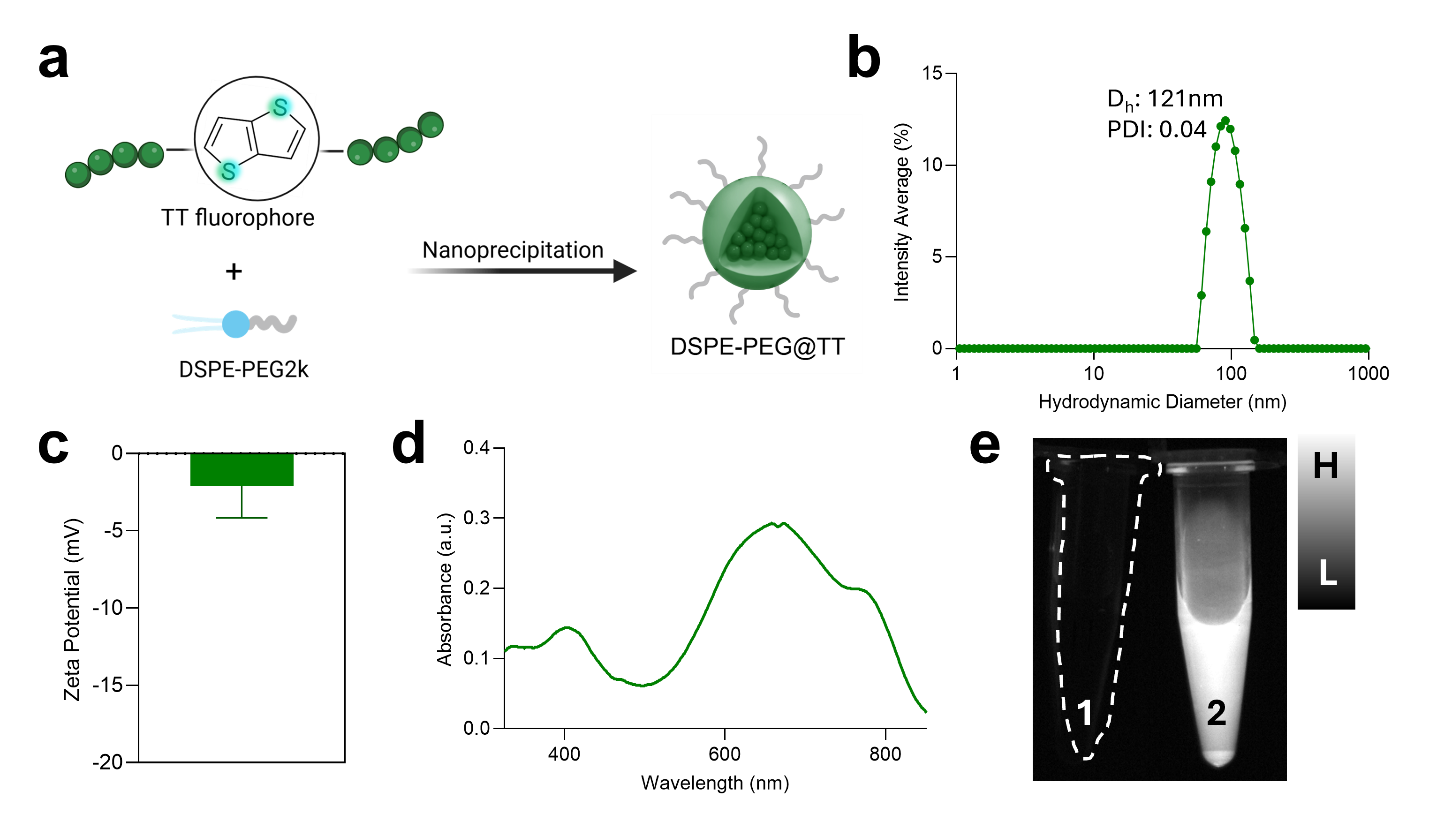

**Figure S7. Physiochemical characterization of DSPE-PEG@TT nanoparticles.** (a) Schematic representation of the preparation of DSPE-PEG@TT. (b) Hydrodynamic diameter and PDI measurements *via* dynamic light scattering. (c) Zeta potential, in mV. (d) UV-Vis-NIR absorbance spectra. (e) IR Vivo fluorescence image of PBS1X (1) and DSPE-PEG@TT nanoparticles (2) with a $\lambda$_ex._ = 808 nm and $\lambda$_em._ = 1000 – 1600 nm. Corresponding scale bar is provided for comparison.

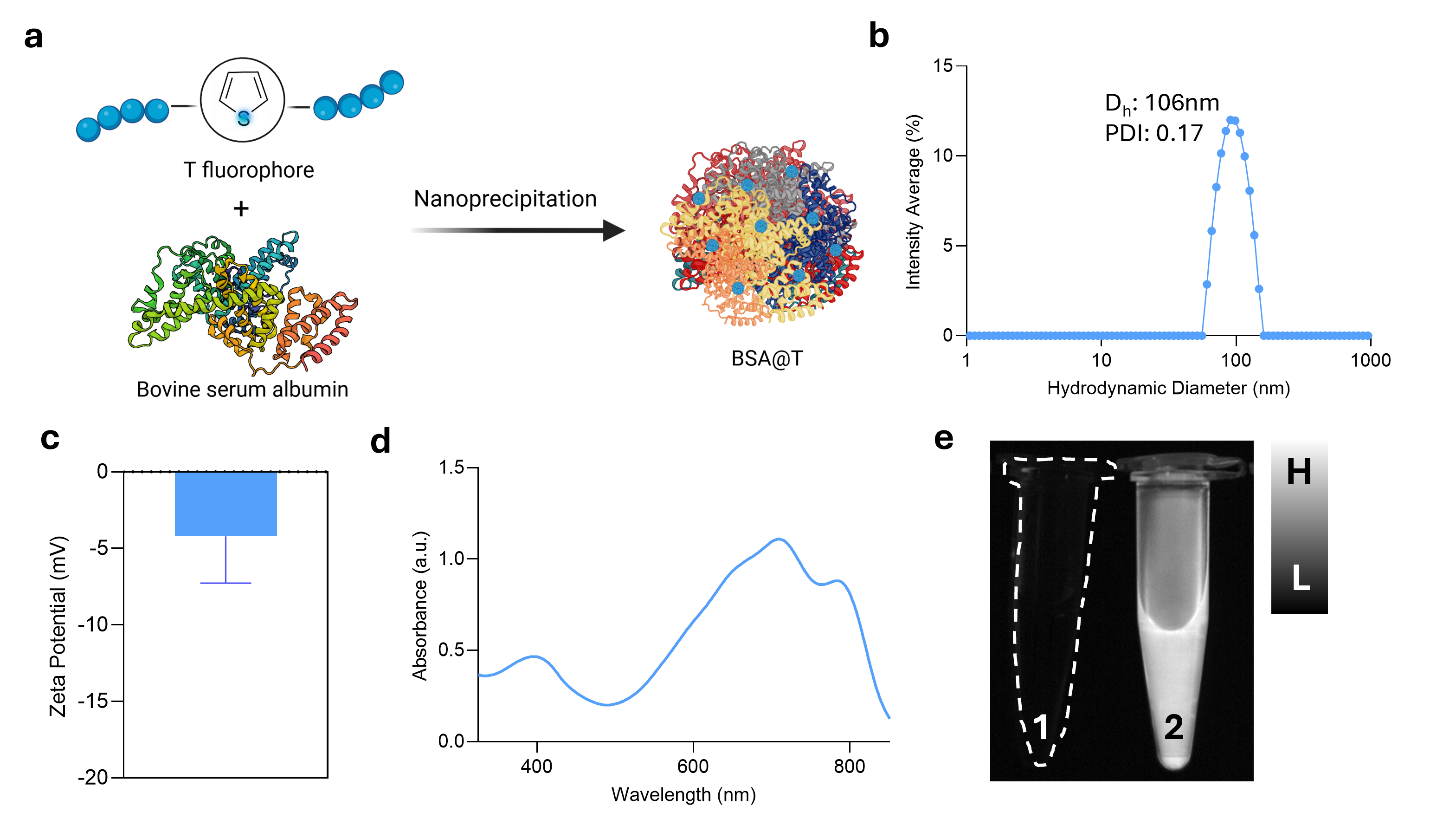

**Figure S8. Physiochemical characterization of BSA@T nanoparticles.** (a) Schematic representation of the preparation of BSA@T. (b) Hydrodynamic diameter and PDI measurements *via* dynamic light scattering. (c) Zeta potential, in mV. (d) UV-Vis-NIR absorbance spectra. (e) IR Vivo fluorescence image of PBS1X (1) and BSA@T nanoparticles (2) with a $\lambda$_ex._ = 808 nm and $\lambda$_em._ = 1000 – 1600 nm. Corresponding scale bar has been shared for comparison.

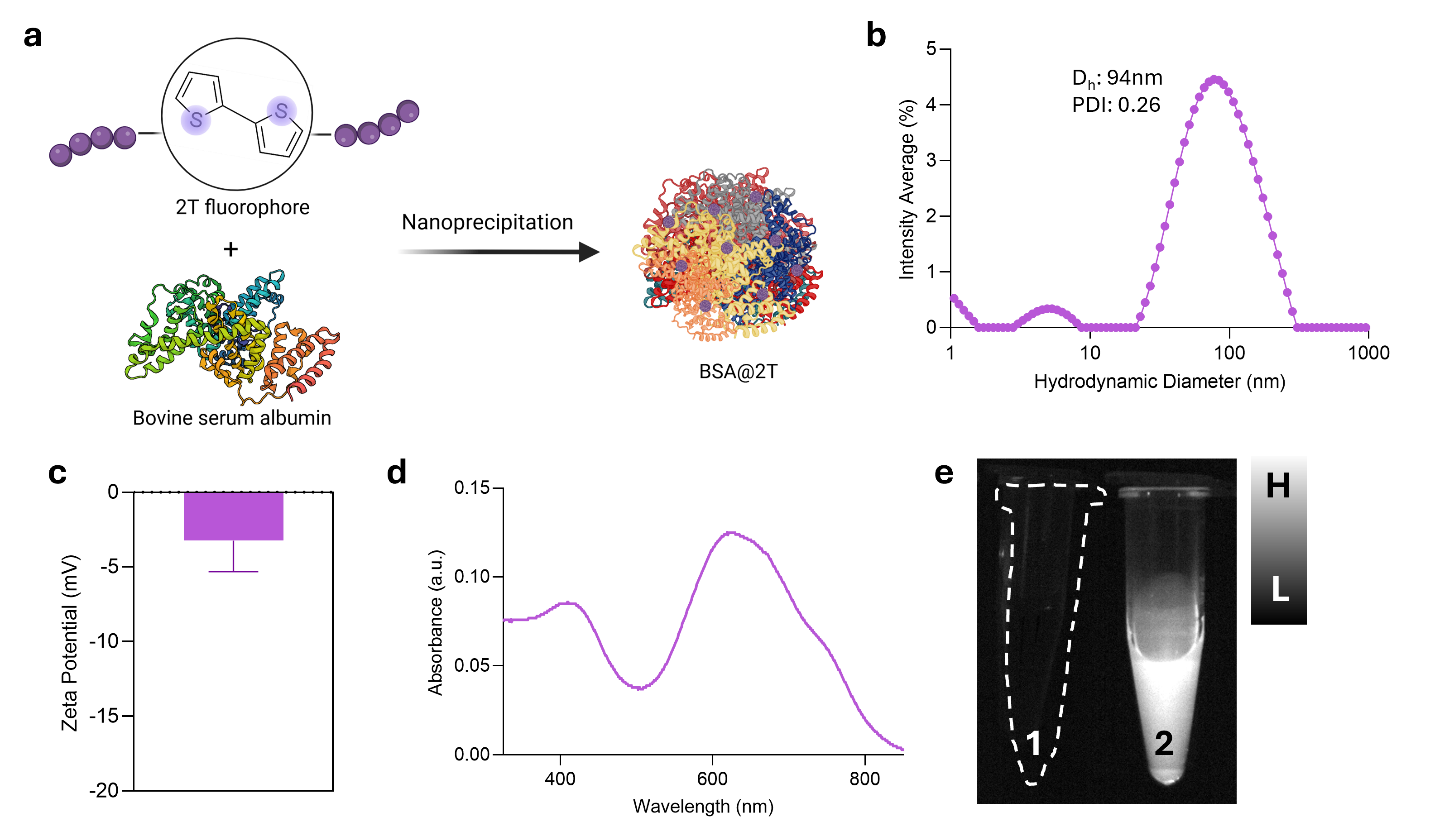

**Figure S9. Physiochemical characterization of BSA@2T nanoparticles.** (a) Schematic representation of the preparation of BSA@2T. (b) Hydrodynamic diameter and PDI measurements *via* dynamic light scattering. (c) Zeta potential, in mV. (d) UV-Vis-NIR absorbance spectra. (e) IR Vivo fluorescence image of PBS1X (1) and BSA@2T nanoparticles (2) with a $\lambda$_ex._ = 808 nm and $\lambda$_em._ = 1000 – 1600 nm. Corresponding scale bar is provided for comparison.

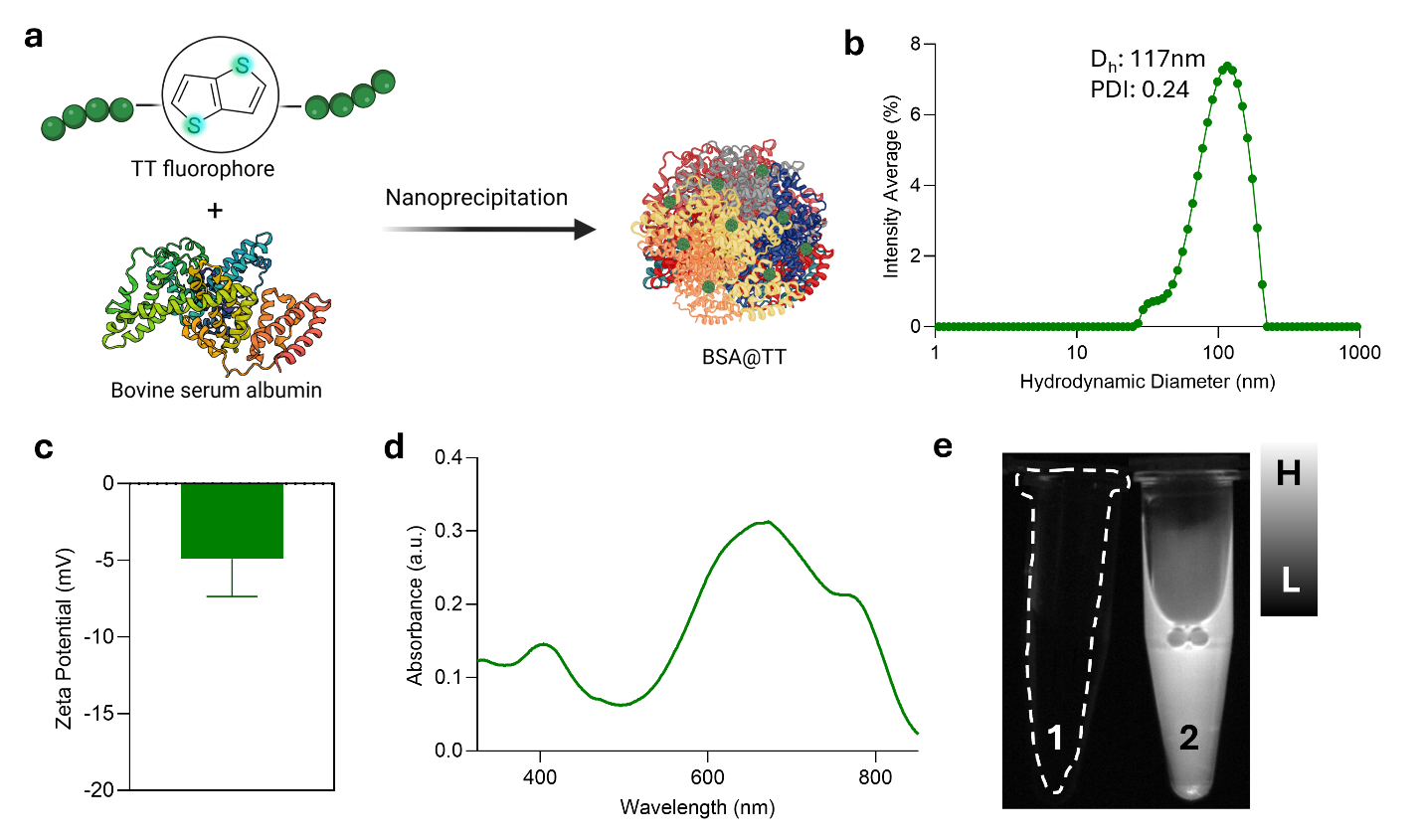

**Figure S10. Physiochemical characterization of BSA@TT nanoparticles.** (a) Schematic representation of the preparation of BSA@TT. (b) Hydrodynamic diameter and PDI measurements *via* dynamic light scattering. (c) Zeta potential, in mV. (d) UV-Vis-NIR absorbance spectra. (e) IR Vivo fluorescence image of PBS1X (1) and BSA@TT nanoparticles (2) with a $\lambda$_ex._ = 808 nm and $\lambda$_em._ = 1000 – 1600 nm. Corresponding scale bar is provided for comparison.

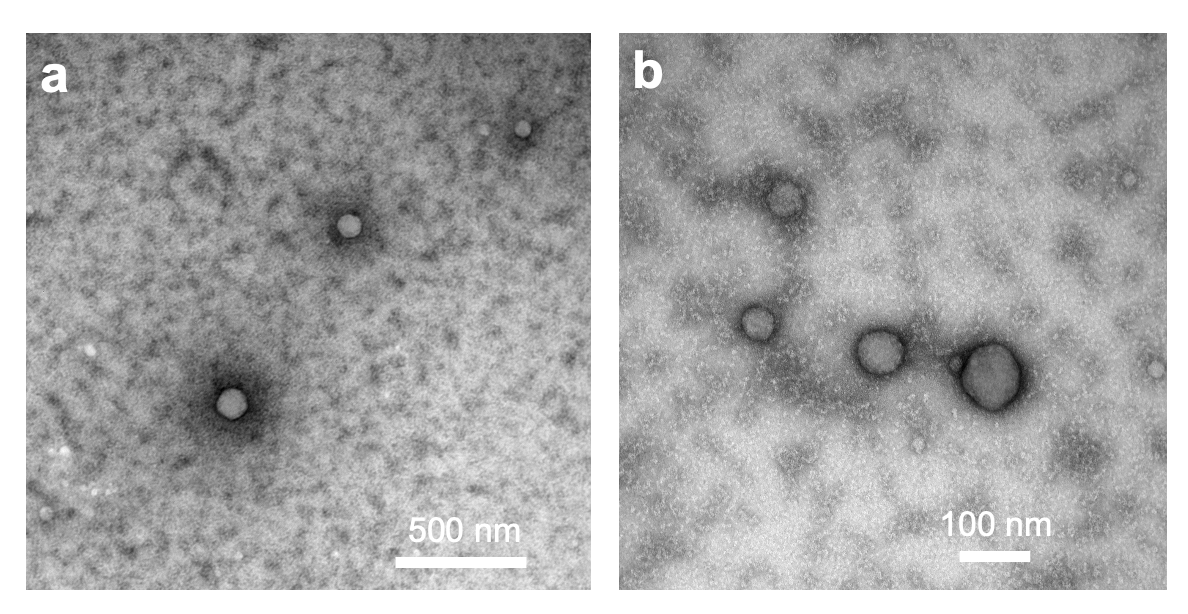

**Figure S11.** (a, b) Transmission electron microscopy images of BSA@TT nanoprobes.

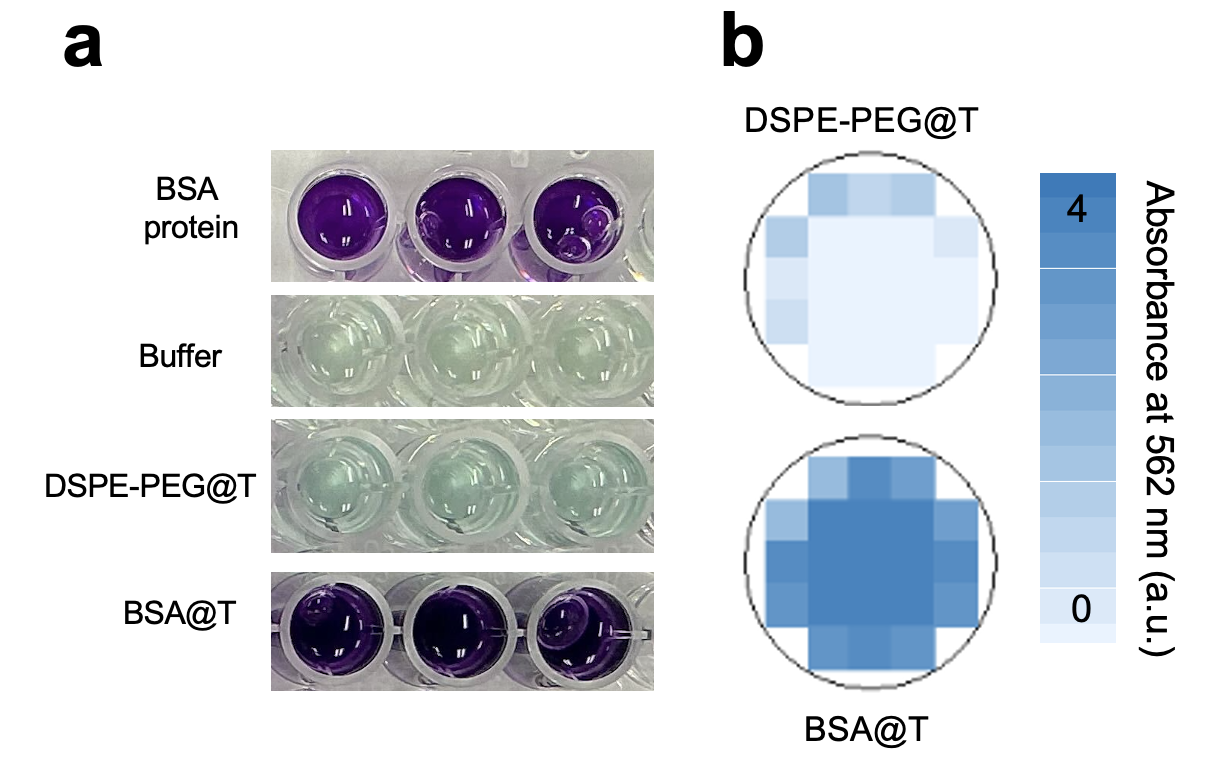

**Figure S12.** Bicinchoninic acid (BCA) assay of nanoprobes measured at 562 nm. Well plate photographs (a) and (b) surface plots of absorbance show strong protein signals for BSA-based formulations (BSA@T, BSA@TT), evidenced by both absorbance peaks and purple coloration. In contrast, DSPE-PEG–coated controls (e.g., DSPE-PEG@TT) show no detectable signal or color change, confirming the absence of protein.

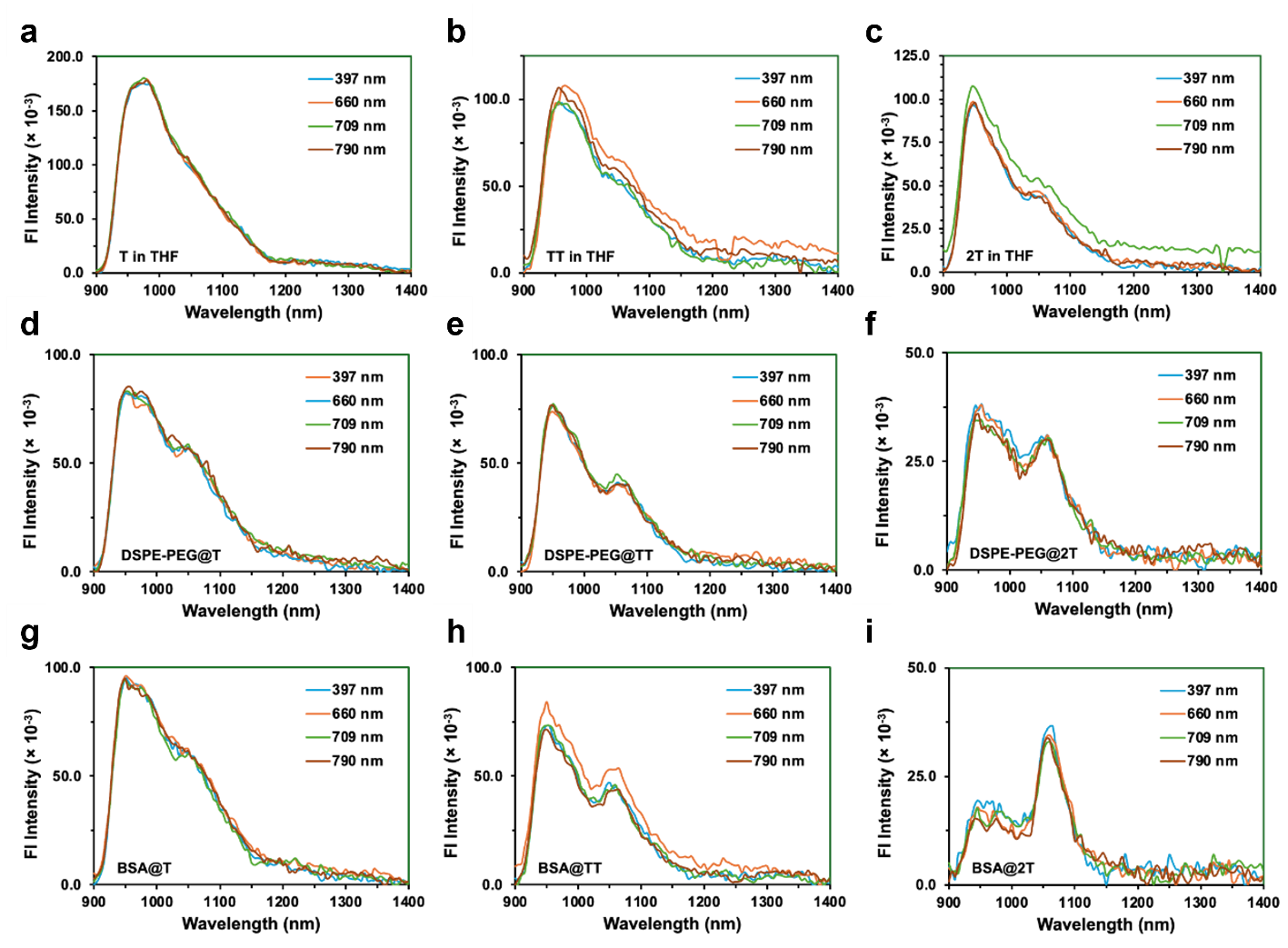

**Figure S13**. Excitation-dependent emission of the SPs (a,b,c), DSPE-PEG NPs (d,e,f) and BSA NPs (g,h,i) at 397, 660, 709, and 790 nm.

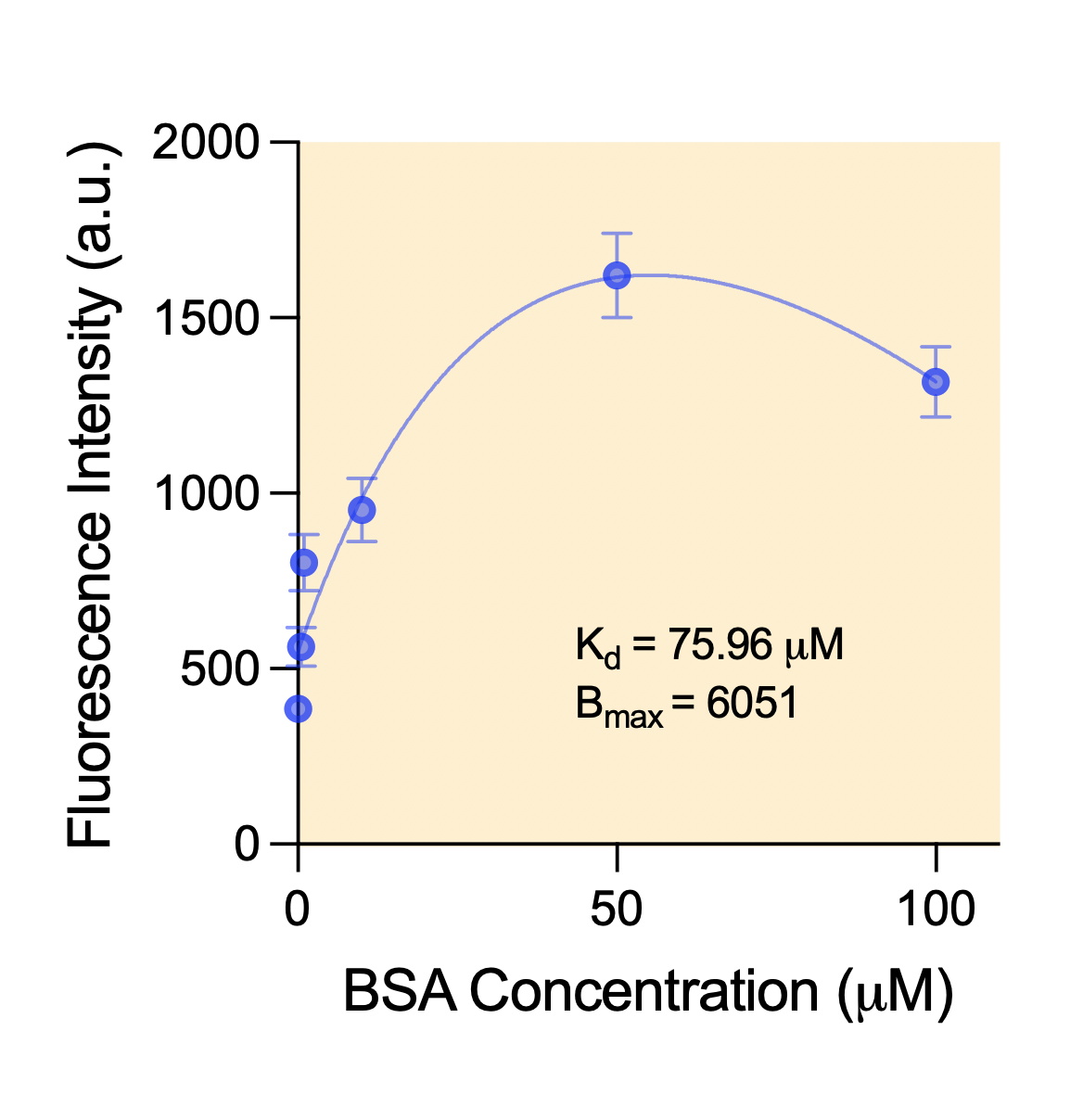

**Figure S14.** Fluorescence titration assay of BSA-TT interaction. Increasing BSA concentrations were incubated with TT polymer, and fluorescence intensity was measured at 900 nm. The binding curve shows saturation behavior, yielding a dissociation constant of *K*_d_ = 75.96 µM and a maximum binding capacity of *B*_max_ = 6051 a.u, confirming specific binding between BSA and TT as predicted by molecular docking.

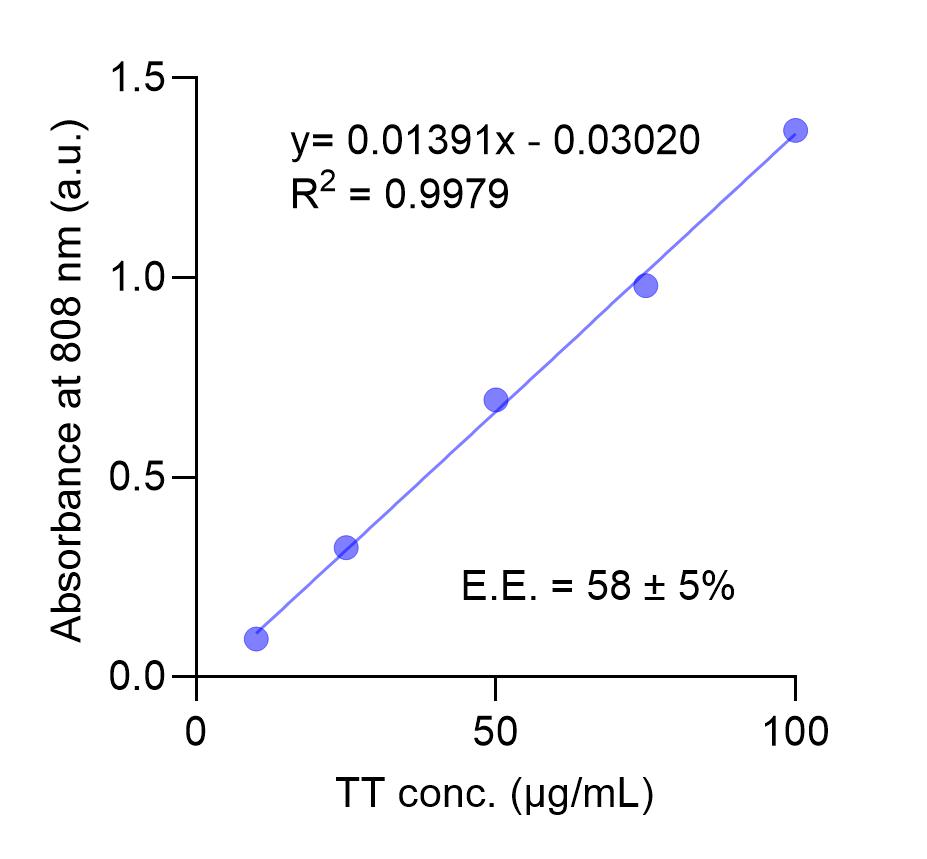

**Figure S15.** UV–Vis calibration and encapsulation efficiency of BSA@TT nanoprobes. (a) Calibration curve generated from TT standards in THF (10, 25, 50, 75, 100 µg/mL), showing linear correlation between absorbance at the maximum wavelength and concentration. (b) Representative UV–Vis spectrum of BSA@TT, with absorbance values converted to TT concentration using the calibration curve. Encapsulation efficiency was calculated as the percentage of TT incorporated into the nanoprobes relative to the initial amount of TT used during formulation.

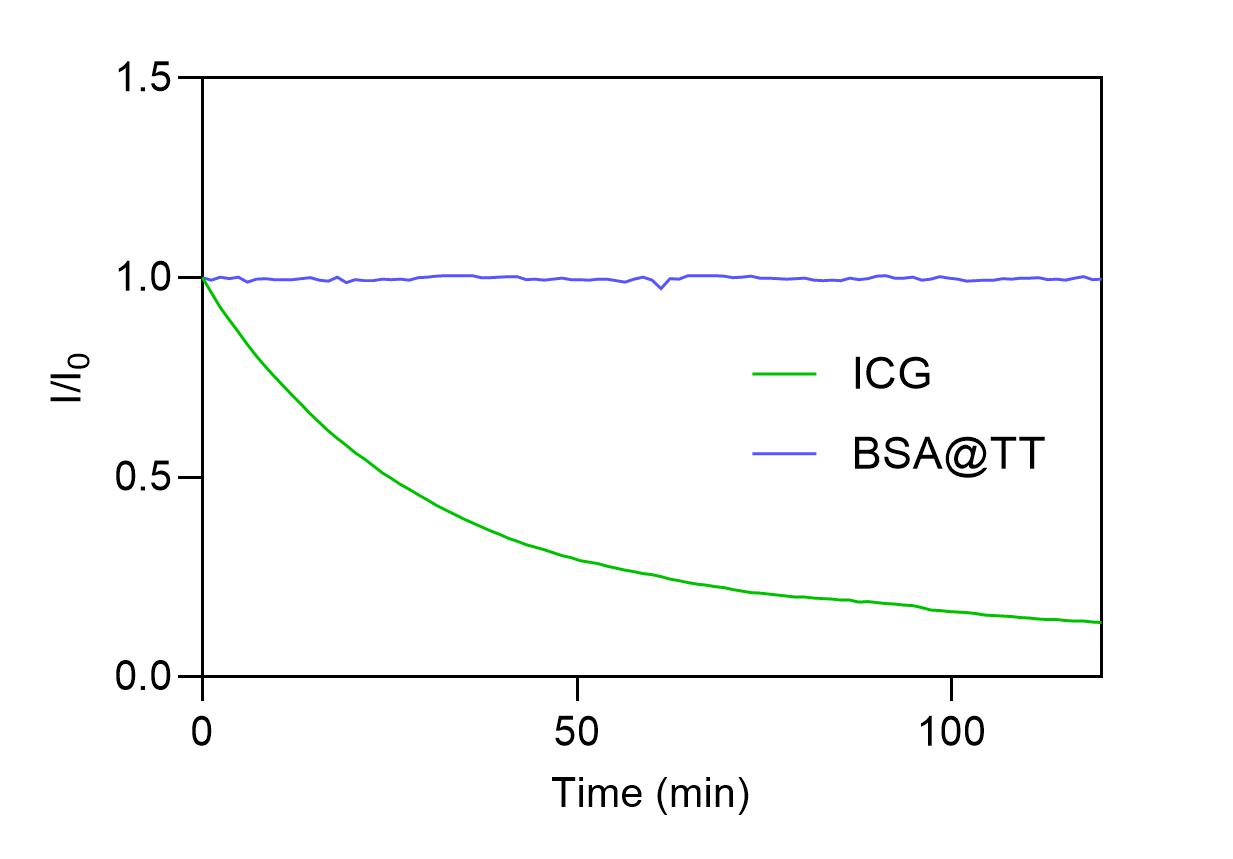

**Figure S16.** Photostability of the BSA@TT and ICG in PBS under continuous irradiation at 808 nm and collection at their respective maximum emission peaks (λ_em,max_).

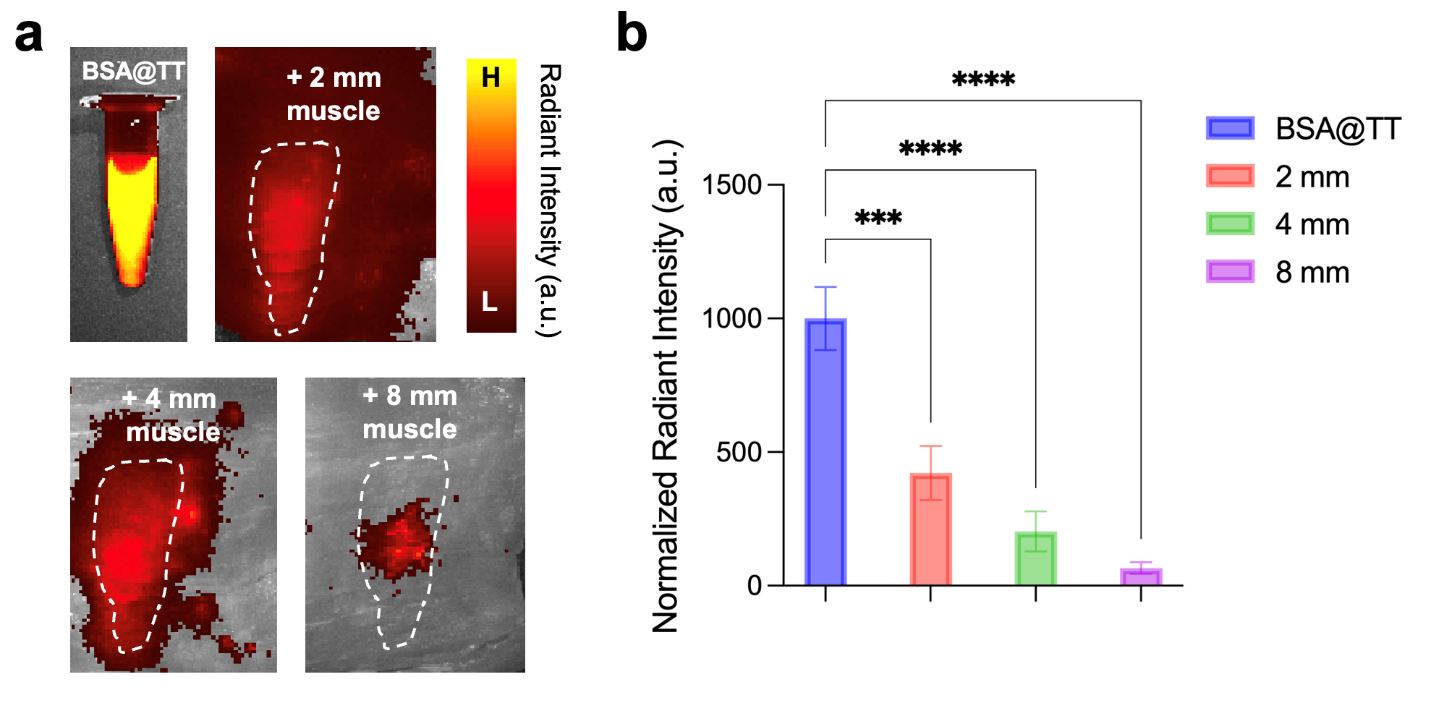

**Figure S17.** Penetration depth of BSA@TT nanoprobes through porcine muscle tissue. (a) Representative NIR-II images of BSA@TT signal with increasing tissue thickness (2, 4, 8 mm). (b) Quantified radiant intensity showing progressive attenuation with depth but detectable signal through 8 mm of muscle. Data are presented as mean ± SD.

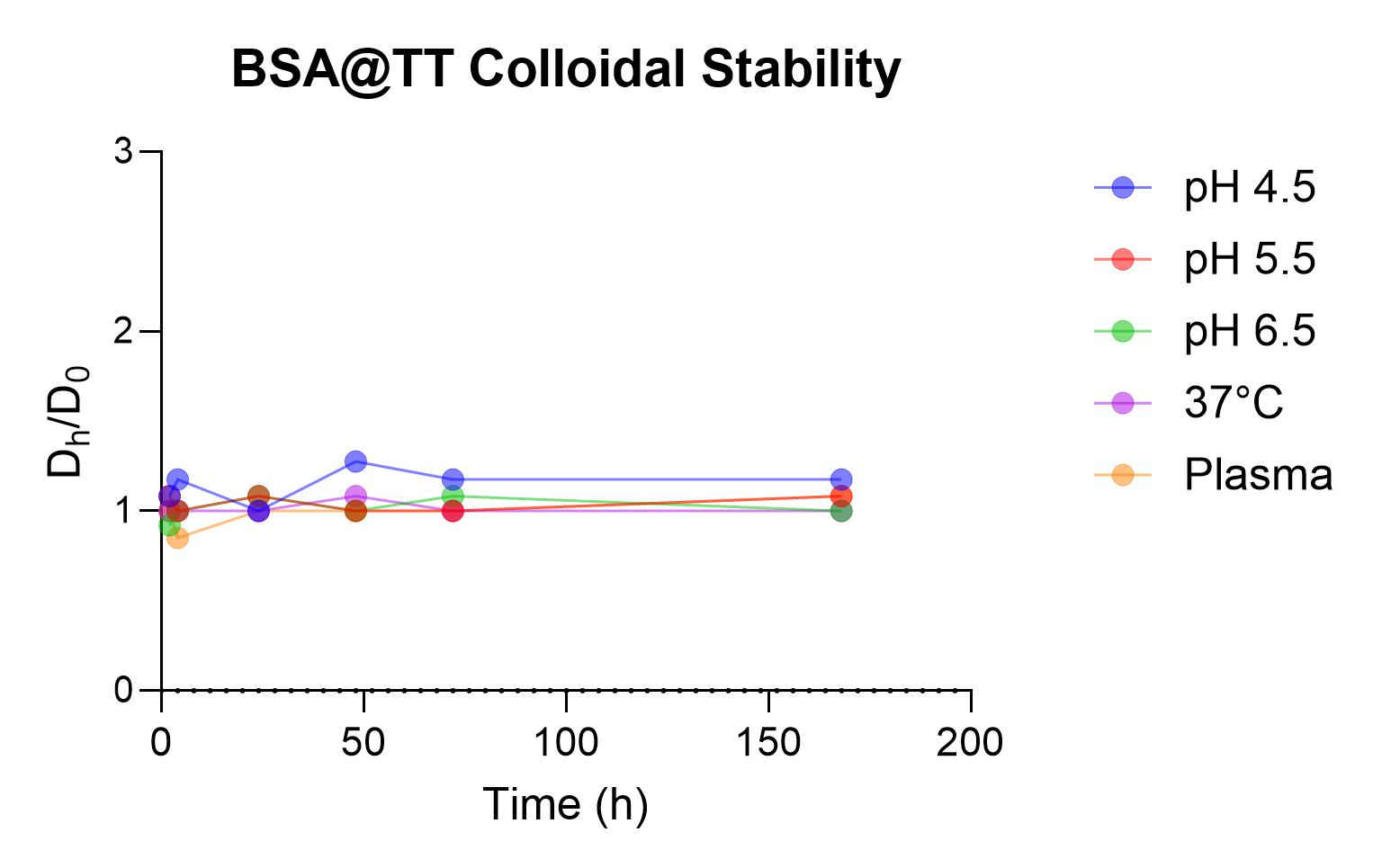

**Figure S18. Colloidal stability of BSA@TT nanoparticles in varying storage and physiologically relevant environments.** Hydrodynamic diameter of BSA@TT NPs when subjected to various pH (4.5, 5.5, and 6.5), 37 ºC, and plasma conditions throughout 1 week.

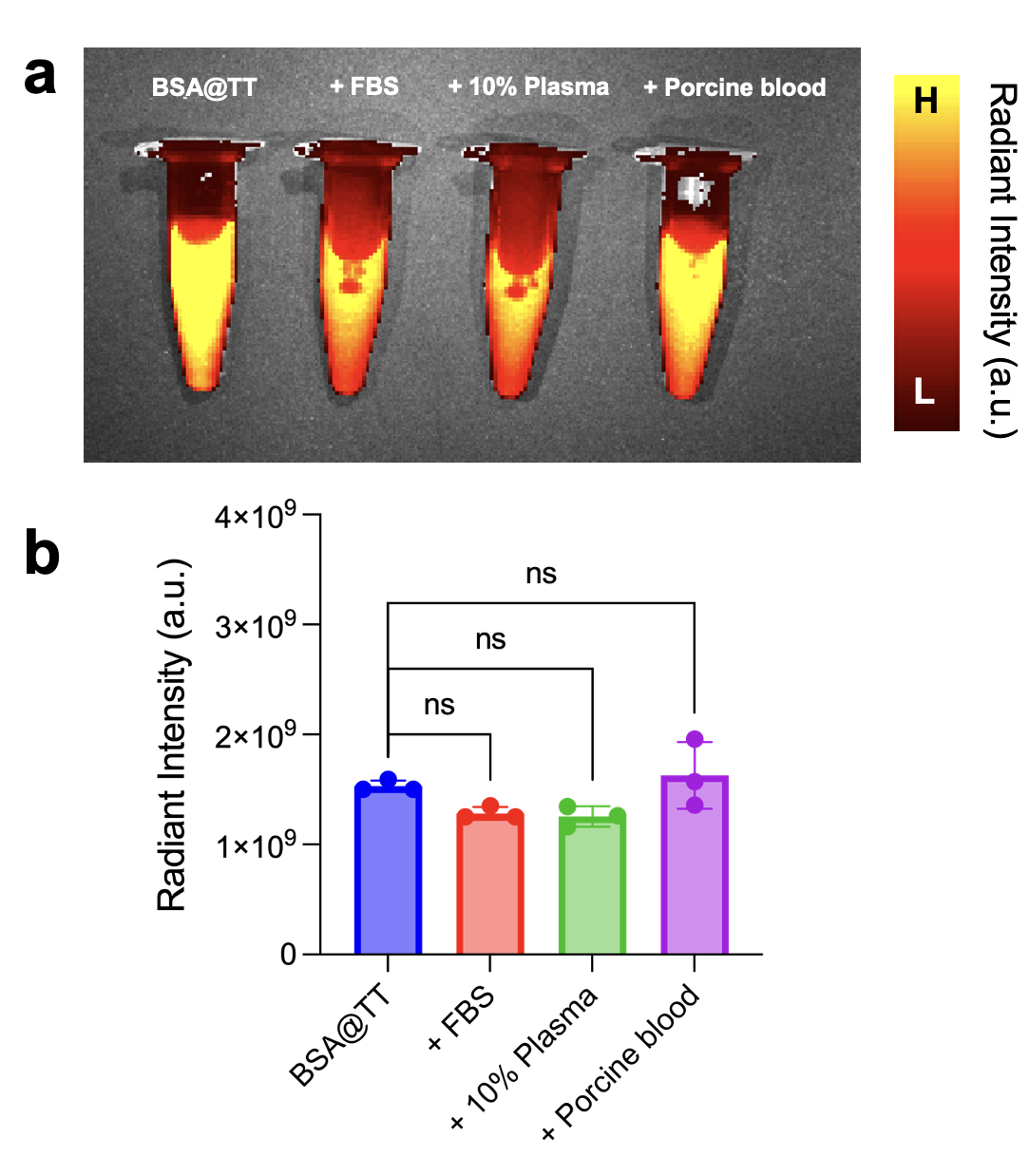

**Figure S19.** Fluorescence stability of BSA@TT nanoprobes in different biological fluids. (a) Representative IVIS images of BSA@TT incubated in PBS, fetal bovine serum (FBS), 10% plasma, and porcine blood. (b) Quantified radiant intensities showed no significant differences across conditions (ns, one-way ANOVA). Data are presented as mean ± SD.

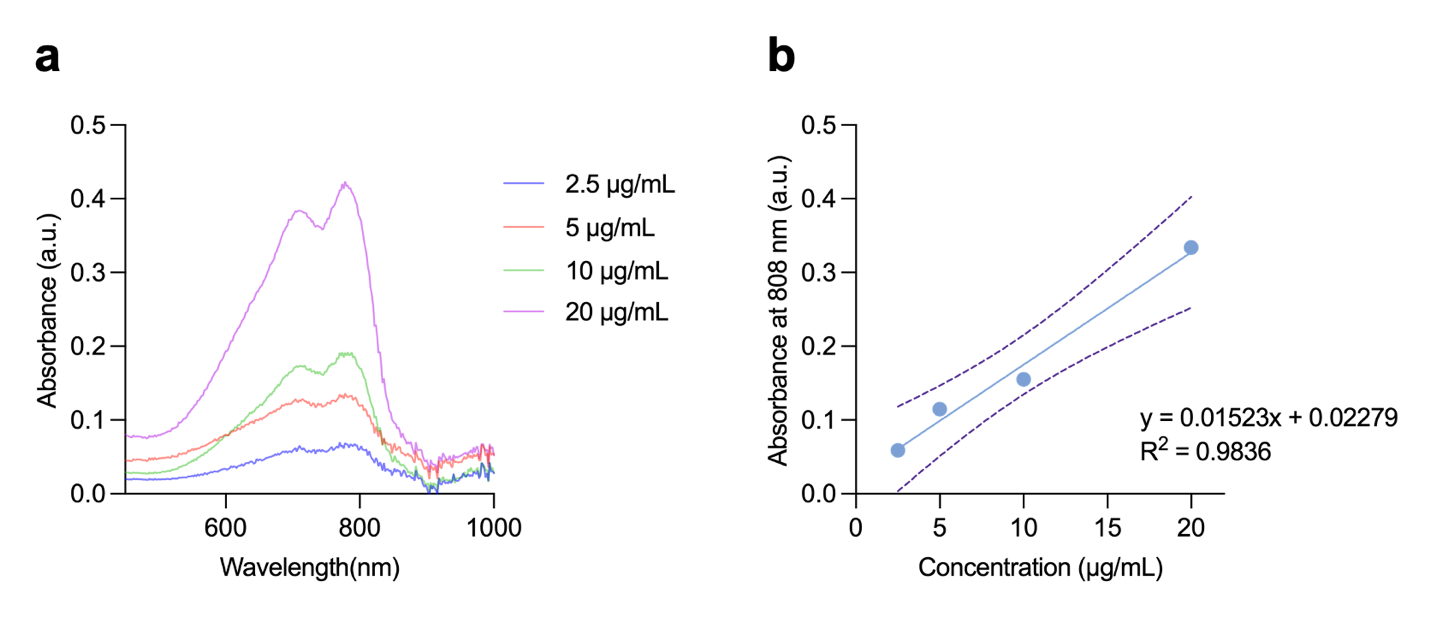

**Figure S20.** Measurement of the mass extinction coefficient of TT in THF. (a) UV–vis–NIR spectra of TT in THF at different concentrations. (b) Absorbance of TT at 808 nm plotted as a function of the concentration of TT in THF. The slope represents the mass extinction coefficient of TT at 808 nm in THF (15.23 L/(g⋅ cm) or 15230 L/mg⋅cm).

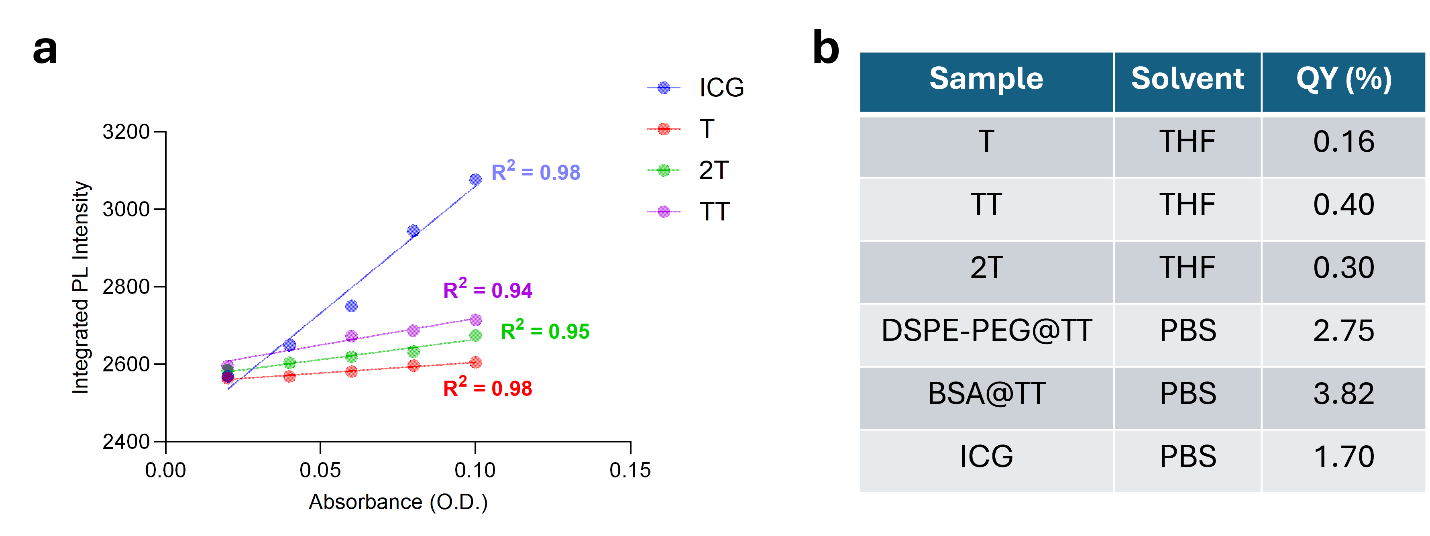

**Figure S21.** (a) Integrated PL intensity versus absorbance plots for ICG, T, 2T, and TT in THF, and encapsulated TT (DSPE-PEG@TT and BSA@TT) in PBS. (b) Table summarizing the quantum yield (QY). Among all tested systems, BSA@TT displays the highest QY (3.82%), compared to DSPE-PEG@TT (2.75%) and TT in THF (0.399%), demonstrating the optical advantages of protein nanoengineering.

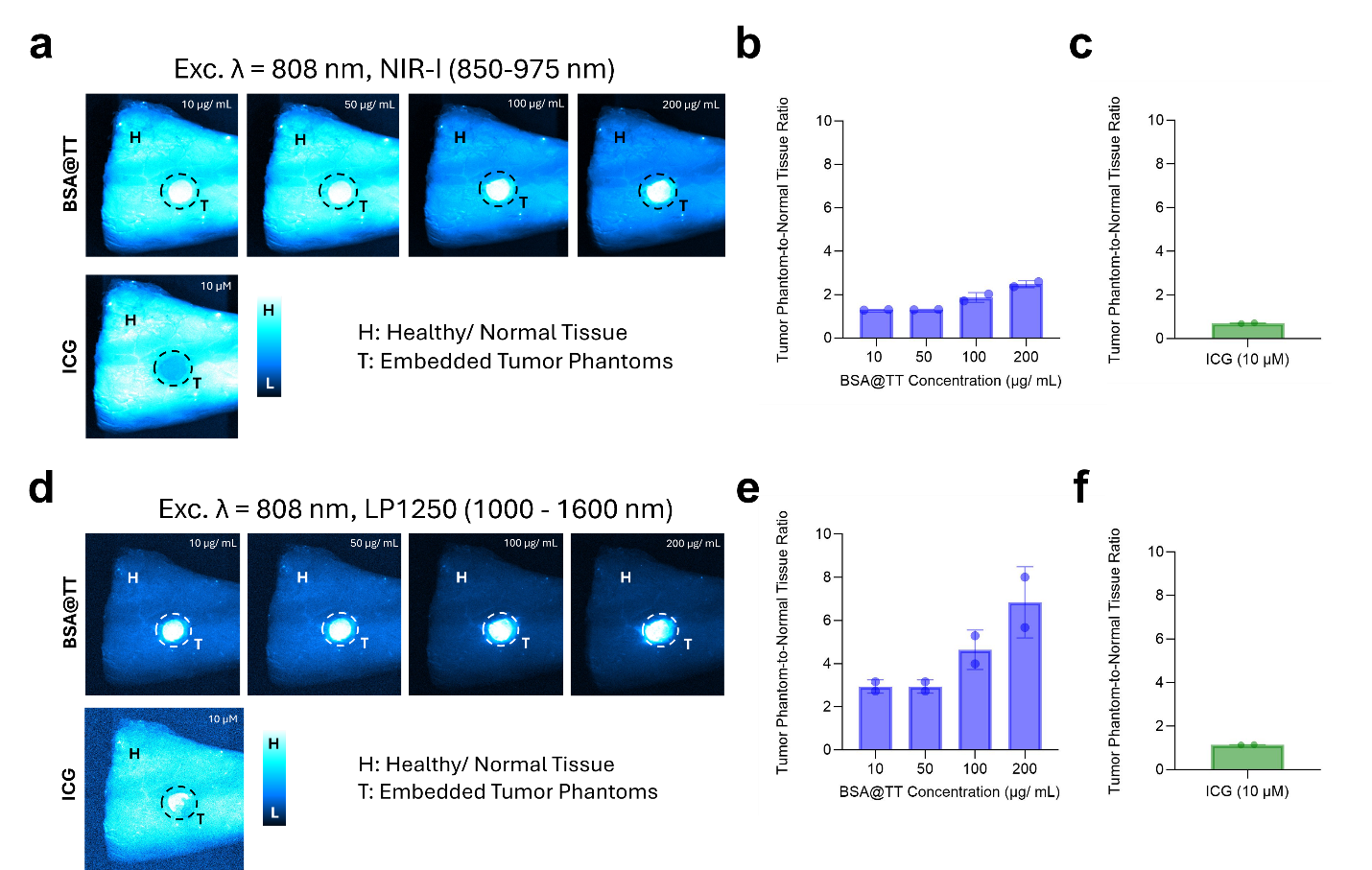

**Figure S22.** **Image-guided surgery simulation using *ex vivo* porcine lung tissue.** IR Vivo fluorescence images with $\lambda$_ex._ = 808 nm of tumor-mimicking phantoms incorporating BSA@TT at increasing concentrations (10, 50, 100, and 200 µg/mL) and ICG, captured within (a) NIR-I and (d) LP1250 emission windows. Tissue discrimination capabilities are reported as tumor phantom-to-normal tissue ratios within the (b), (c) NIR-I and (e), (f) LP1250 emission windows.

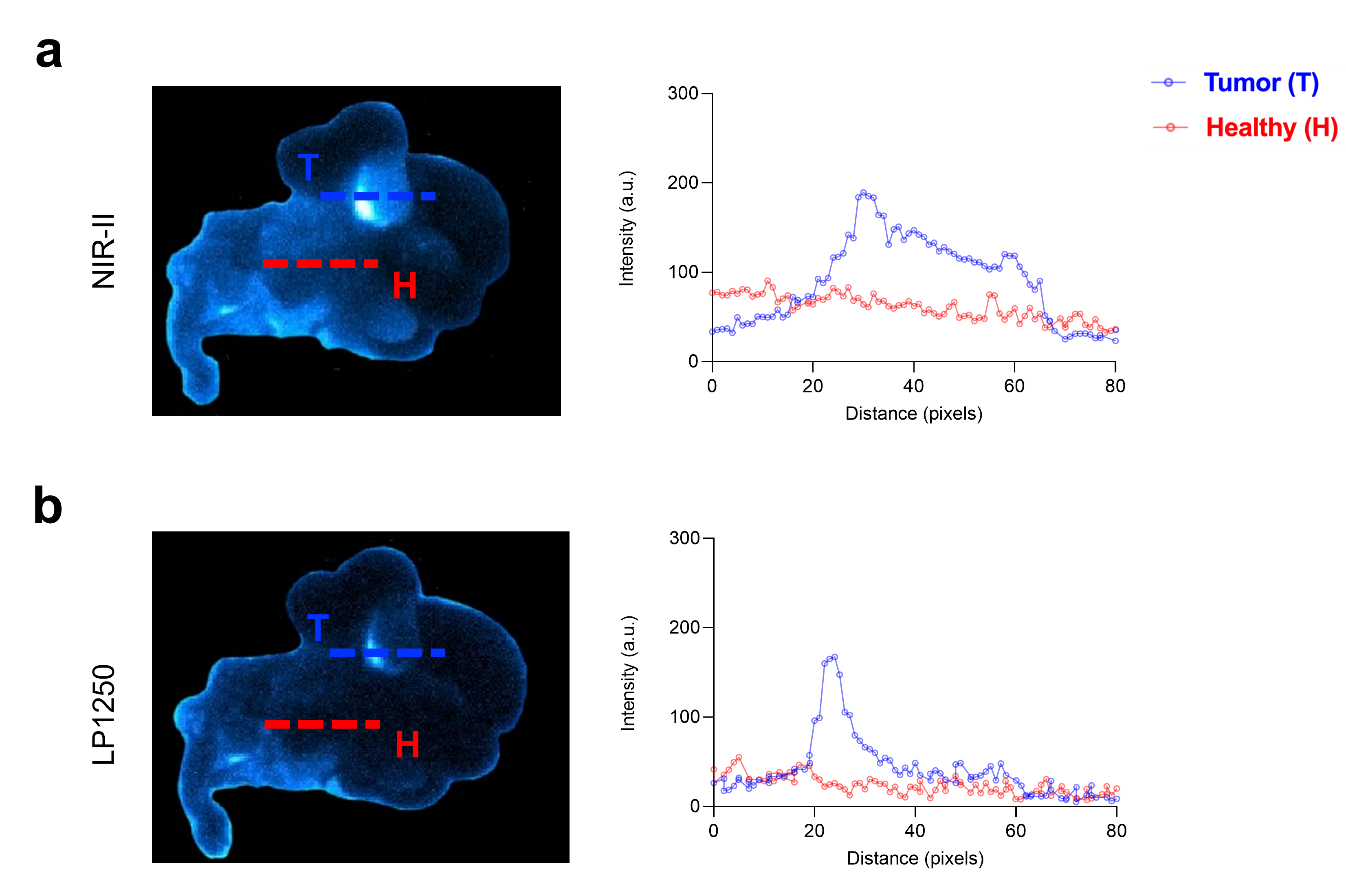

**Figure S23.** **Discriminating the “tumor tissue” from surrounding healthy tissues in *ex vivo* porcine ovary models with BSA@TT NPs.** Representative NIR-II fluorescence images and line intensity graphs (n=3 injections) of a BSA@TT-filled porcine ovary follicle at *λ*_ex_: 808 nm with (a) *λ*_em_: 1000 – 1600 nm and (b) *λ*_em_: 1250 – 1600 nm.

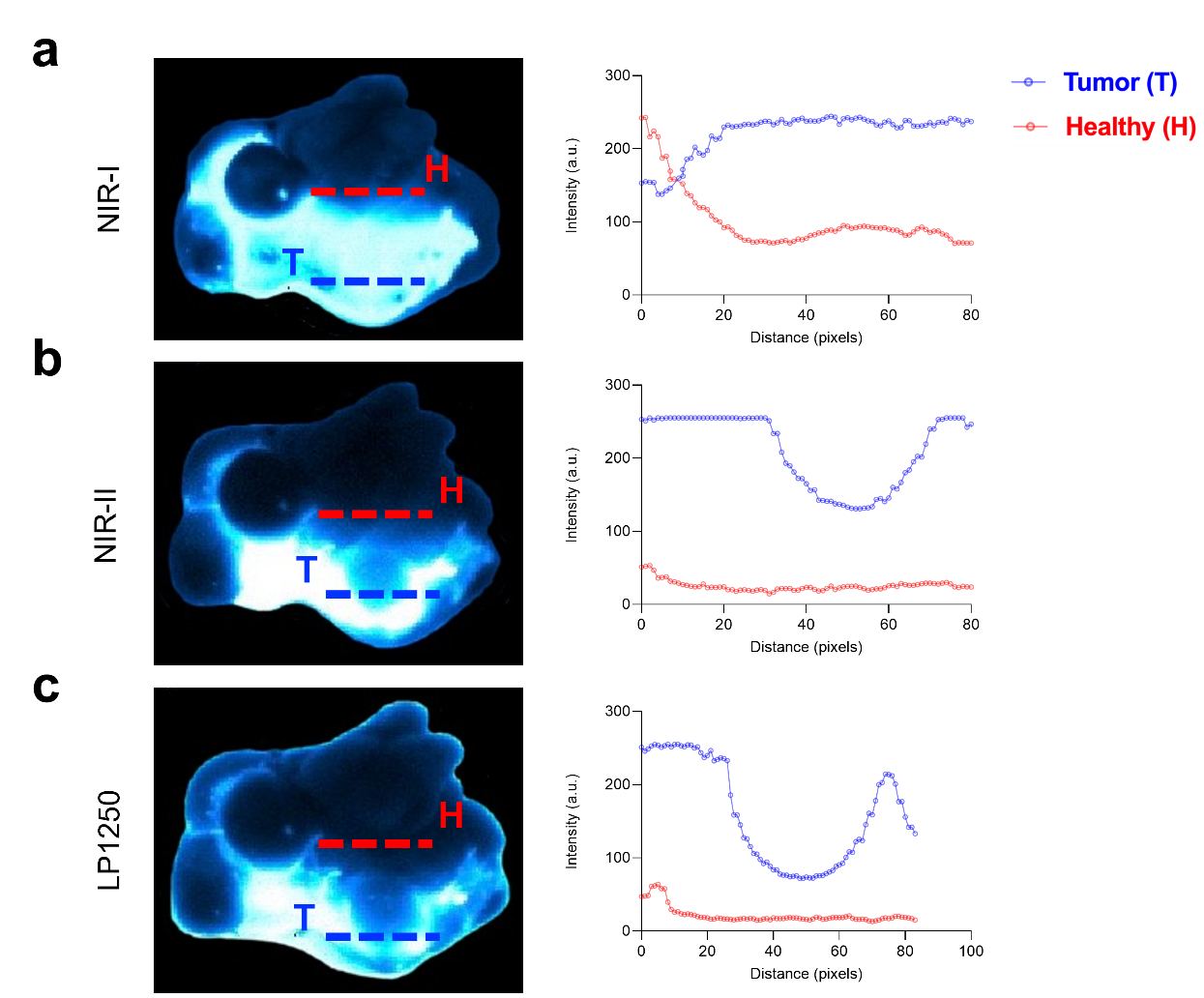

**Figure S24.** **Discriminating “tumor tissue” from surrounding healthy tissue in *ex vivo* porcine ovary models with ICG.** Representative NIR-II fluorescence images and line intensity graphs (n=3 injections) of an ICG-filled porcine ovary follicle at *λ*_ex_: 808 nm with (a) *λ*_em_: 850 – 975 nm, (b) *λ*_em_: 1000 – 1600 nm, and (c) *λ*_em_: 1250 – 1600 nm.

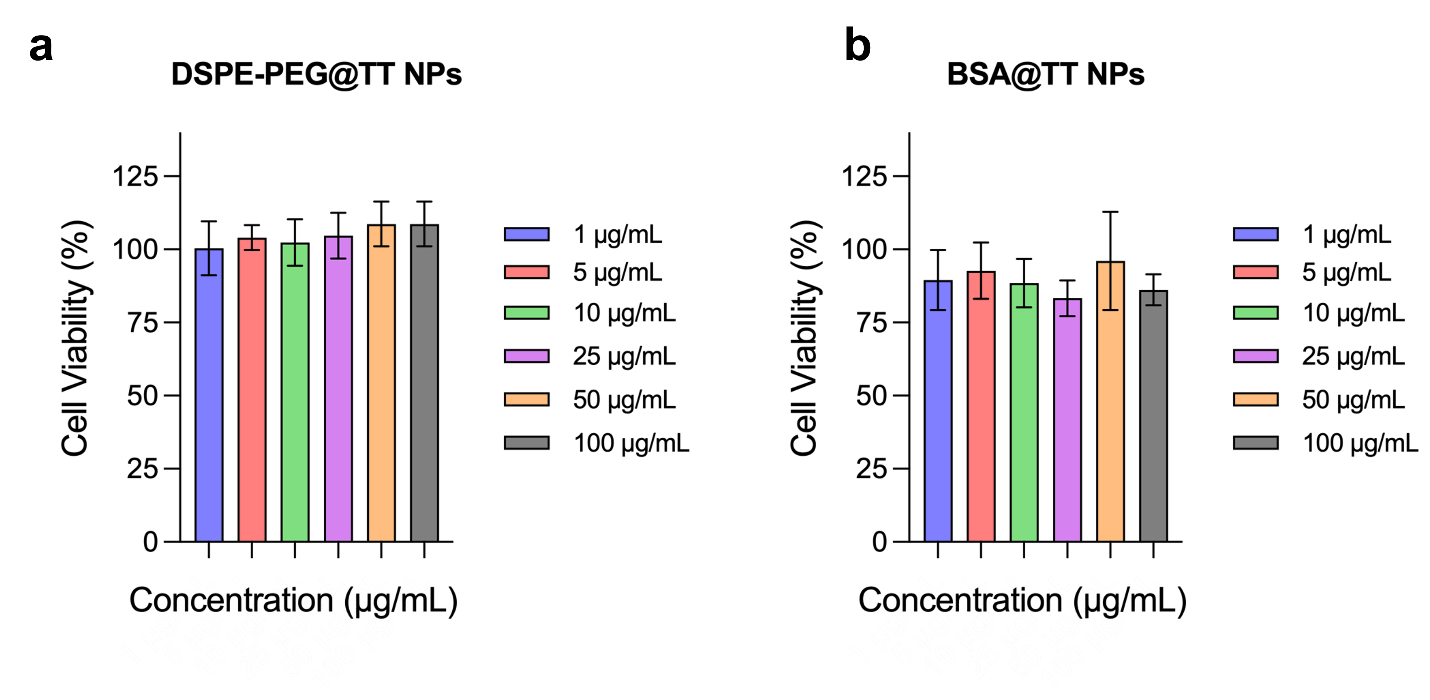

**Figure S25.** **Assessing Cytotoxicity *via* MTT Assay.** Cell viability of OVCAR8 cells following incubation with (a) DSPE-PEG@ TT and (b) BSA@TT nanoparticles, respectively, at increasing concentrations: 1, 5, 10, 25, 50, and 100 µg/mL. Experiments were performed after incubation for 48 h at these different concentrations.

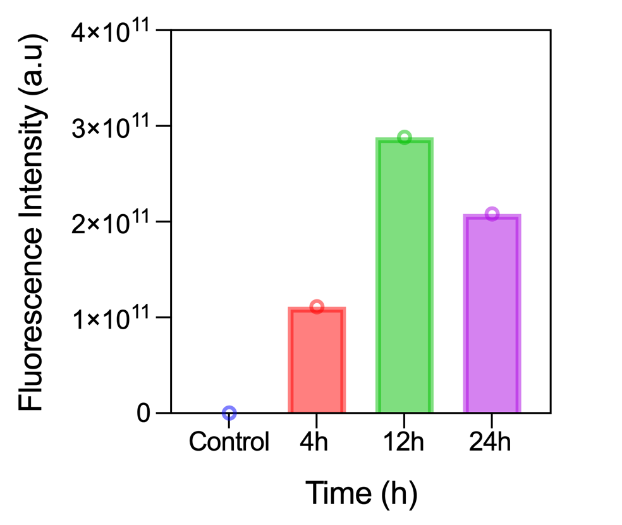

**Figure S26. Cellular uptake of BSA@TT nanoprobes in OVCAR8 cells.** Quantification of fluorescence intensity indicates time-dependent internalization of BSA@TT nanoprobes at 4, 12, and 24 hours.

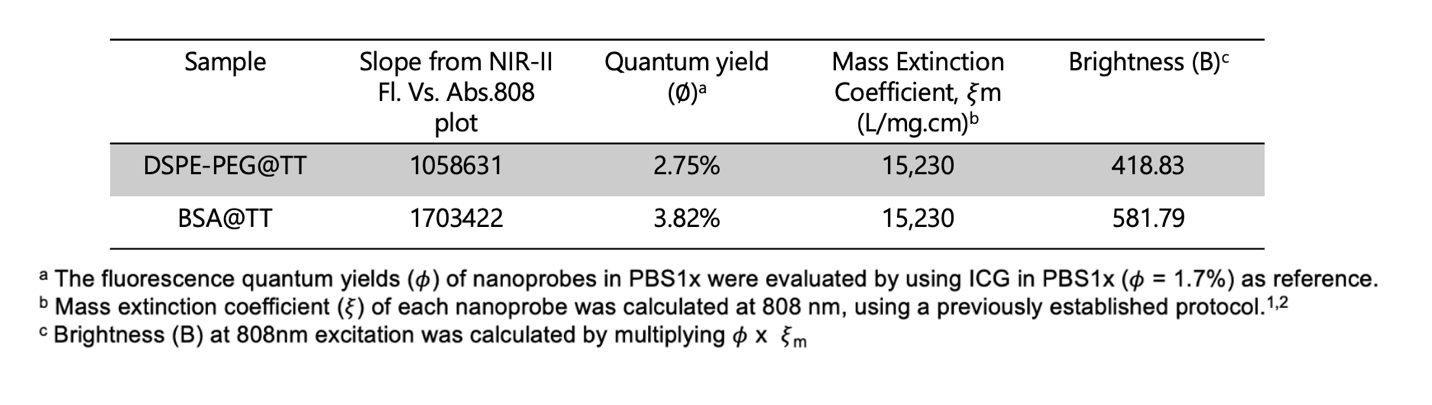

**Table S1.** Optical properties evaluation of BSA@TT and DSPE-PEG@TT nanoprobes.

**References.**

(1) Zhou, H.; Lu, Z.; Zhang, Y.; Li, M.; Xue, D.; Zhang, D.; Liu, J.; Li, L.; Qian, J.; Huang, W. Simultaneous Enhancement of the Long-Wavelength NIR-II Brightness and Photothermal Performance of Semiconducting Polymer Nanoparticles. *ACS Applied Materials & Interfaces* **2022**, *14* (7), 8705-8717. DOI: 10.1021/acsami.1c20722.

(2) Zhang, X.; Yang, Y.; Kang, T.; Wang, J.; Yang, G.; Yang, Y.; Lin, X.; Wang, L.; Li, K.; Liu, J.; et al. NIR-II Absorbing Semiconducting Polymer-Triggered Gene-Directed Enzyme Prodrug Therapy for Cancer Treatment. *Small* **2021**, *17* (23), 2100501. DOI: doi.org/10.1002/smll.202100501.
